# Decoding G0 somatic mutants through deep phenotyping and mosaic pattern analysis in the zebrafish skeleton

**DOI:** 10.1101/466185

**Authors:** Claire J. Watson, Adrian T. Monstad-Rios, Rehaan M. Bhimani, Charlotte Gistelinck, Andy Willaert, Paul Coucke, Yi-Hsiang Hsu, Ronald Y. Kwon

**Affiliations:** Department of Orthopaedics and Sports Medicine, University of Washington, Seattle, Washington, USA; Institute for Stem Cell and Regenerative Medicine, University of Washington, Seattle, Washington, USA; Department of Mechanical Engineering, University of Washington, Seattle, Washington, USA; Center for Medical Genetics Ghent, Ghent University, Ghent, Belgium; Hebrew SeniorLife Institute for Aging Research, Boston, Massachusetts, USA; Harvard Medical School, Boston, Massachusetts, USA

**Keywords:** CRISPR, crispant, zebrafish, bone, osteoblast, phenomics, Osteogenesis Imperfecta, brittle bone disease, osteoporosis, *bmp1a*, *plod2*, *wnt16*

## Abstract

Genetic mosaicism manifests as spatially variable phenotypes, whose detection and interpretation remains challenging. This study identifies biological factors influencing spatial phenotypic patterns in the skeletons of somatic mutant zebrafish, and tests methods for their analysis using deep phenotyping. We explore characteristics of loss-of-function clusters in the skeleton of CRISPR-edited G0 ("crispant") zebrafish, and identify a distinctive size distribution shown to arise from clonal fragmentation and merger events. Using microCT-based phenomics, we describe diverse phenotypic manifestations in somatic mutants for genes implicated in monogenic (*plod2* and *bmp1a*) and polygenic (*wnt16*) bone diseases, each showing convergence with germline mutant phenomes. Finally, we describe statistical frameworks for phenomic analysis which confers heightened sensitivity in discriminating somatic mutant populations, and quantifies spatial phenotypic variation. Our studies provide strategies for decoding spatially variable phenotypes which, paired with CRISPR-based screens, can identify genes contributing to skeletal disease.

## INTRODUCTION

The past decade has seen a steady rise in genomics, fueled by advances in next-generation sequencing that have improved the speed and efficiency by which we can sequence genomes. Analogous to innovations in genome sequencing, advances in phenomics—i.e., in depth phenotyping at a large number of anatomical sites—can advance our understanding of how genetic variation influences phenotype, including biology that involves relationships between phenotypes across the whole organism (1). A particular example wherein phenomic profiling would lend such insight is in the context of genetic mosaicism: the presence of cells with multiple distinct genotypes constituting the organism on the whole. Mosaicism can arise naturally through errors in DNA replication, or intentionally through genetic manipulation. This genetic heterogeneity results in a hallmark of mosaicism—site-to-site phenotypic variability—which makes identifying gene-to-phenotype relationships challenging. In animal models, somatic mutations form the basis for rapid-throughput G0 screens, prototypes for which are rapidly increasing following the advent of CRISPR (Clustered-Regularly Interspaced Short Palindromic Repeats)-based gene editing (2, 3). In humans, chromosomal mosaicism in embryos is quite common (4, 5), its role in disease may be prevalent and underappreciated (6, 7), and at the most fundamental level, it is suggested that every complex, multi-cellular organism is likely to harbor at least some, if even very low-level, somatic mosaicism (8, 9). In the context of disease, mosaic individuals can be affected by mutations that clinically manifest in their offspring, and it has been suggested that phenotypic patterns can inform the timing of mutagenesis, and thus, the likelihood of mutations in the germline and that can be passed onto progeny (9). Deep phenotypic profiling at a number of anatomical sites may help decode somatic mutant phenotypes, which are important in both experimental and clinical settings.

A prime instance of experimental biology which necessitates decoding somatic mutant phenotypes is in a rapid-throughput genetic screen. When complexed with a targeting guide RNA (gRNA), the bacterial Cas9 enzyme will create a double strand break at a genome-specific location determined by complementary sequence in the gRNA. Errors in the endogenous non-homologous end joining (NHEJ) repair mechanism lead to a high rate of insertions and deletions (indels) at the cut site, often leading to frameshift mutations and loss of gene product function. Several prototypes for rapid CRISPR-based reverse genetic screens have been developed in which phenotyping is performed directly in G0 founders (10, 11). This increases throughput by alleviating the time and resources needed to breed mutant alleles to homozygosity. Such approaches may also be useful for animal models that require longer durations to reach sexual maturity or have long gestational intervals which render multiple rounds of breeding to homozygosity unfeasible. However, creating universal loss-of-function is challenging. For instance, when administered a single gRNA, 1/3 of indels are expected to be in-frame. Thus, less than half ([2/3]^2^ = 4/9) of cells are expected to have bi-allelic out-of-frame mutations (10). While the use of multiple guides to redundantly target the same gene can increase the proportion of bi-allelic out-of-frame mutations, this may also increase toxicity, and variable penetrance of null phenotypes is still prevalent (11). Microhomology-mediated end joining (MMEJ) is a promising direction to enrich somatic mutations for a predictable out-of-frame allele (12). However, imperfect editing efficiencies and some degree of mutated allelic mosaicism are still expected following MMEJ. Finally, an advantage of G0 screens is that they enable multiplexed gene knockdown, which can be used to study epistatic interactions of genes that are tightly linked, or knockdown of clusters of genes with functional redundancy (10, 13, 14). Yet, mutation efficiency often decreases with the number genes that are multiplexed, due to the reduction in Cas9:gRNAs per gene. Due to the lower fidelity in detecting somatic mutant phenotypes, prototypical screens have mostly focused on severely dysmorphic phenotypes (11), or phenotypes whose spatial variations are easily observable (10).

Extracting biological information from somatic mutant phenotypes remains challenging for several reasons. One source of difficulty is our lack of understanding of quantitative phenotypic variation arising from mosaicism, and how best to analyze it. For spatially distributed organs (e.g., bone, skin, nerves, blood vessels), mosaicism can manifest as relatively uniform phenotypes reminiscent of a generalized condition, or alternating patterns of affected and unaffected body segments (15). Much of our knowledge of the phenotypic consequences of mosaicism has been derived from easily observable traits where spatial variations are readily discernible (16). As such, our ability to discern phenotypic manifestations of mosaicism for complex traits remains relatively limited (16). While technologies for phenomic profiling in vertebrates are increasing, workflows have mostly been defined in germline mutants (1, 17, 18) or animals subjected to systemic drug exposure (19). Different analytical methods may be needed for somatic mutants, where the specific set of altered measures, acquired from different anatomical locations, may be different from animal-to-animal. Moreover, most statistical methods established for-omics data do not account for spatial relationships between measures. Assimilation of such spatial information is essential to detect differences in spatial patterns among different somatic mutant groups. Robust phenomic workflows for somatic mutant analysis could help realize the full potential of CRISPR-based rapid throughput biology, and serve as a prototype to better understand clinical manifestations of mosaicism in human diseases (20, 21).

Another source of difficulty is our limited understanding of biological factors that influence phenotypic expressivity in mosaic individuals. It is broadly accepted that spatial phenotypic patterns are dependent on the proliferation of mutant cells and their translocation to different sites. Lineage tracing of clonal populations from embryonic to adult zebrafish has provided a wealth of knowledge on clonal abundance in a variety of tissues, and suggests that, in most cases, a few clonal progenitors account for a majority of cells comprising the resulting tissue type (22). Yet, how these clones distribute spatially within and across tissues remains unknown, and is a critical piece of information needed to interpret somatic mutant phenotypes. Spatial phenotypic patterns can also be influenced by the function of the mutated gene itself. For example, the degree to which phenotypic patterns mirror patterns of mosaicism likely depends on whether the gene acts cell autonomously or non-cell autonomously (20, 21). While this implies that phenotypic patterns in mosaic individuals encode information related to gene function that is not present in germline mutants, instances of phenotypic pattern recognition in somatic mutant analysis are not well described.

Here, we identify patterns of mosaicism in the skeletons of CRISPR-edited somatic zebrafish, and relate these clonal distributions to phenotypes in mosaic models of brittle bone diseases. We employ a microCT-based workflow enabling profiling of hundreds of measures per fish (1) to characterize quantitative phenotypic variation, and perform experimentally-informed simulations to test methods for discerning and interpreting somatic mutant populations. Finally, we provide a case study for these methods by identifying and characterizing a novel zebrafish axial skeletal mutant whose target gene has been linked to genetic risk for osteoporosis.

## RESULTS

### CRISPR-based gene editing results in clusters of cells with loss-of-function

To examine patterns of loss-of-function in the skeleton, *sp7:EGFP* (23) embryos were injected with Cas9:guide RNA (gRNA) ribonucleoprotein complexes (RNPs) targeting the fluorescent transgene. This enabled loss-of-function mutations in EGFP to be visualized as loss-of-fluorescence in *sp7*+ (*osterix*+) osteoblasts. Fish injected with 10μM RNPs were examined for functional EGFP loss at 10-12dpf, a stage when the larvae are still transparent and many skeletal elements are present. Regions of loss-of-fluorescence were observed in virtually all formed skeletal elements (Fig 1A). This included bones of the craniofacial skeleton (opercle, interopercle, subopercle, dentary, pharyngeal arches, branchiostegal rays, cleithrum), median fin rays (ventral, caudal), hypurals, and the spine (neural and haemal arches, centra, Weberian apparatus). While penetrance of loss-of-fluorescence was high, expressivity was variable in regard to the composition of bony elements in each animal that exhibited regions of fluorescence loss, as well as the size and number of such regions within each bony element.

**Fig 1.**
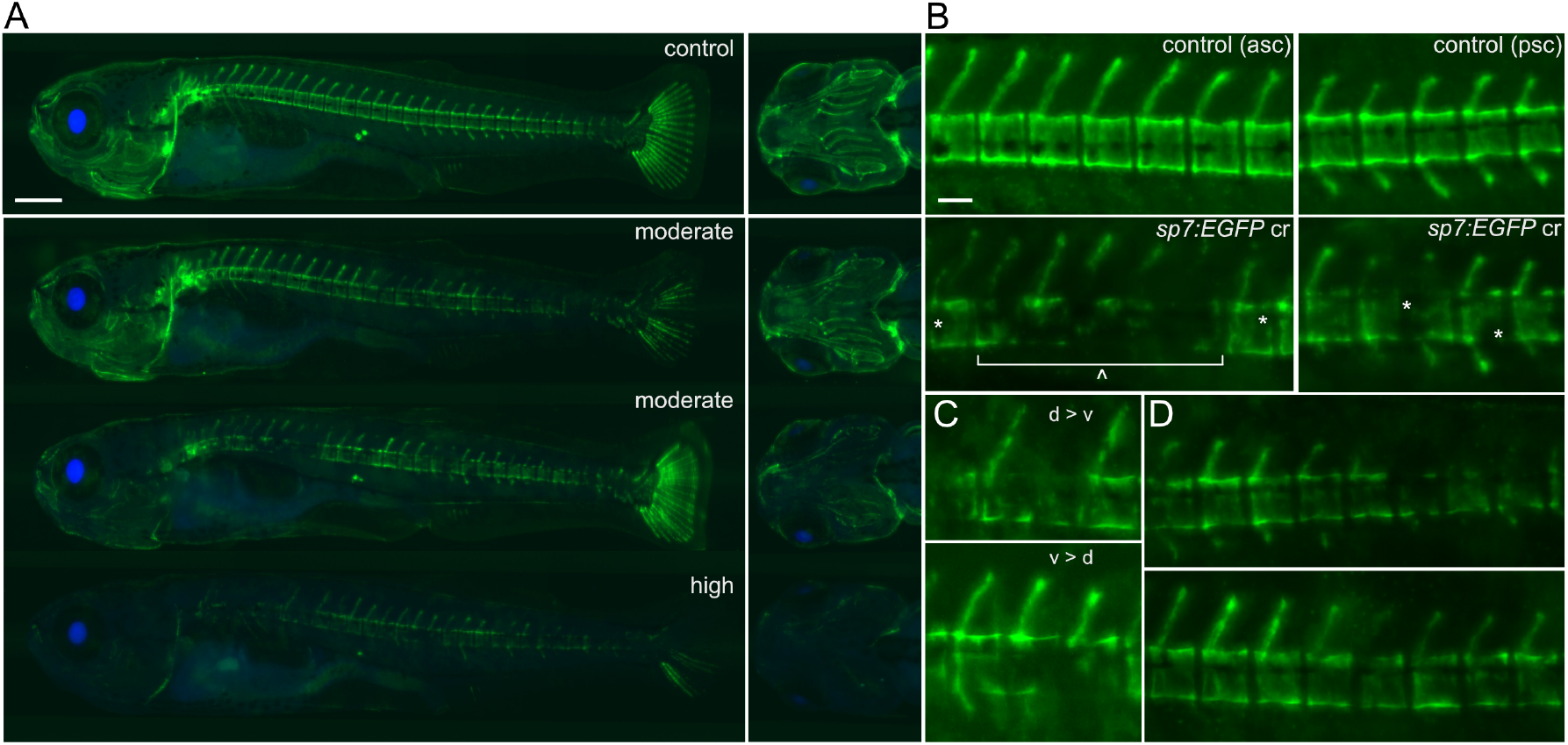
Somatic mutations in the *sp7:EGFP* transgene show mosaic loss of fluorescence. (A) Larval *sp7:EGFP* transgenic fish show relatively uniform EGFP expression in the skeleton of control fish (top). Somatic mutants with either a moderate (middle) or a high (bottom) degree of loss-of-fluorescence display a wide and varied range of loss-of-fluorescence cluster distributions, including in the craniofacial skeleton, the axial skeleton, and in the caudal fin rays. All fish presented are 10 dpf. Scale bar = 500 µm. (B) Loss-of-fluorescence occurs on both the macroscale (spanning multiple vertebral bodies, denoted by a caret) and the microscale (contained within a vertebral body, denoted by asterisks) compared to controls. Notice the distinction between opacity due to developing pigmentation in the controls (top) compared to loss of fluorescence in somatic mutants (bottom). asc, anterior spinal column; psc, posterior spinal column. (C) On multiple instances, loss-of-fluorescence in somatic mutants was stratified along the dorso-ventral axis of the centrum, with loss occurring preferentially on the dorsal side (top) or ventral side (bottom). (D) EGFP expression (or lack thereof) occurring on the dorsal side of the centrum in somatic mutants often corresponded to expression in the neural arch and spine in both the anterior (top) and posterior (bottom) spinal column. Scale bar for (B-D) = 100 µm.

Loss-of-fluorescence regions are composed of clusters of cells with loss-of-function mutations: cells comprising each cluster represent a single clone, or multiple clones that merged at an earlier point in development (we are unable to distinguish these two possibilities). Within individual vertebrae, we often observed multiple, contiguous centra with complete or partial loss-of-fluorescence, and which were flanked by at least one centra with no loss-of-fluorescence. This resulted in two distinguishable types of cell clusters: “microscale” clusters confined within single vertebrae, and “macroscale” clusters spanning contiguous vertebrae (Fig 1B). Inspection at higher magnification revealed that some centra exhibited loss-of-fluorescence in ventral, but not dorsal, regions (or vice versa). This dorso-ventral stratification could at times be seen across contiguous centra (Fig 1C), potentially due to these bodies’ shared clonal partners. In regard to the neural arches, loss-of-fluorescence often appeared to be associated with loss-of-fluorescence in the centrum of the same vertebral body (Fig 1D). Because many bones retained partial or complete expression of the transgene, this suggested that, on the whole, individual bony elements are not explicitly derived from single clonal populations, and cannot be evaluated as independent functional or non-functional units.

### A common distribution underlies the sizes of loss-of-function clusters in bones of distinct developmental lineages, and in animals with different mutation efficiencies

While some aspects of loss-of-fluorescence patterns appeared to be non-random, patterns from fish-to-fish were unpredictable, suggesting stochastic forces were an important etiological factor. Models of clonal population dynamics in fluorescence-based cell lineage tracing studies have demonstrated that while different factors can contribute to cluster size distributions during tissue growth, over time, contributions from random clonal merger and fragmentation (Fig 2A) become dominant over those from cell behaviors specified by developmental programs (e.g., cell division or loss)(24). As a consequence, cluster size distributions across diverse developmental processes often exhibit the same characteristic distribution once cluster sizes in each individual are normalized by the average cluster size in that individual (24). This distribution has the form:

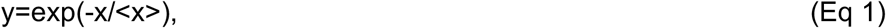

where x is cluster size, and <x> is the mean of × (24).

**Fig 2.**
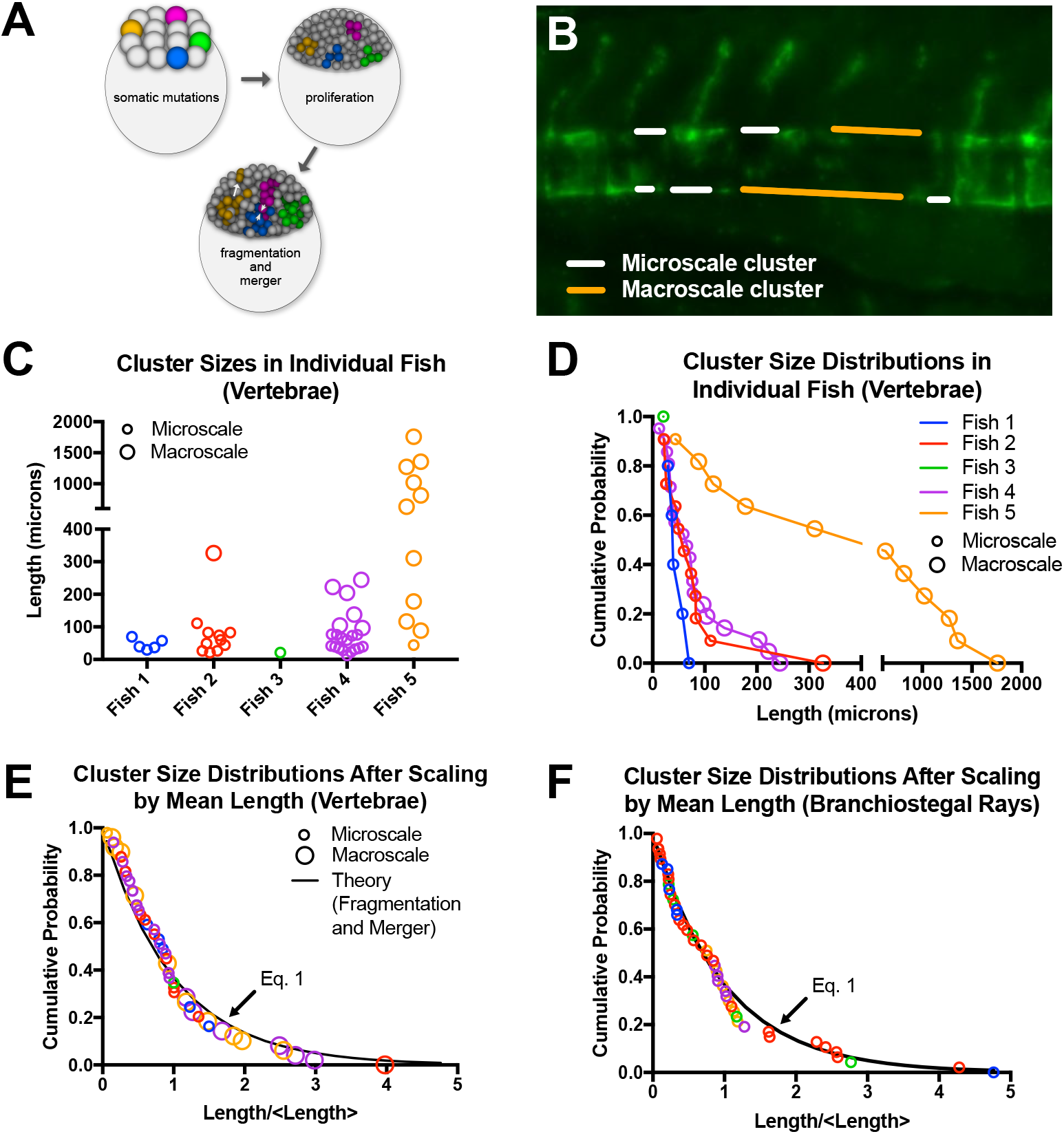
A common distribution underlies the sizes of loss-of-function clusters in bones of distinct developmental lineages, and in animals with different mutation efficiencies. (A) Schematic demonstrating clonal fragmentation and merger events. (B) Schematic demonstrating tracing of microscale (white) and macroscale (orange) loss-of-fluorescence clusters on the dorsal and ventral centrum surfaces. (C) Cluster sizes in vertebrae of individual fish. Note the differences in the number and sizes of clusters, indicative of differences in extent of loss-of-function in each fish. (D) Vertebral cluster size distributions in individual fish, using data from panel (C). (E) Vertebral cluster size distributions in individual fish, when normalized by the mean length in each fish. Color mapping is same as for panels C and D. Note the collapse of data onto a common distribution, which overlaps with Eq. 1. (F) Cluster size distributions for loss-of-function clusters in the branchiostegal rays. Color mapping is same as for panels C and D.

Fluorescently-labeled cell clusters in cell lineage tracing studies and loss-of-fluorescence cell clusters in somatic mutants share commonalities in their physical origins (postzygotic mutations) and interpretation (clusters may be comprised of a single clonal population, or multiple clones that merged earlier in development). As such, we hypothesized that loss-of-function clusters in somatic animals would also exhibit universality in their size distributions described by Eq 1. To test this, we manually traced regions of complete loss-of-fluorescence (defined as a region with no detectable fluorescence above background, and/or markedly reduced signal compared to adjacent bony structures, see “Methods”) on the dorsal and ventral aspects in the centra in each animal (Fig 2B). Regions confined to a single vertebrae were annotated as “microscale clusters”; regions spanning contiguous vertebrae were annotated as “macroscale clusters”. Individual fish exhibited variable numbers and sizes of loss-of-function clusters (Fig 2C), as well as different compositions of microscale versus macroscale clusters. This variability manifested as distinct distributions of loss-of-function cluster sizes in each fish (Fig 2D). However, when normalized by average cluster size in each animal, the cluster size distributions in each animal collapsed to the distribution in Eq 1 for both microscale and macroscale clusters (Fig 2E). We hypothesized that this distribution would also describe loss-of-function cluster sizes in bones of a different developmental lineage. We quantified loss-of-function cluster size distributions within the branchiostegal rays of the craniofacial skeleton, which unlike the somite-derived vertebral column, derives from neural crest (25). Consistent with our hypothesis, a similar data collapse was observed (Fig 2F). These studies demonstrate that loss-of-function cluster sizes in bones of distinct developmental lineages, and in animals with different loss-of-function efficiencies, can be described by a single distribution (Eq 1); the origin of which derives from numerical convergence behaviors associated with clonal fragmentation and merger events.

### Somatic mutants for plod2 and bmp1a exhibit differences in variability in phenotypic expressivity

To better understand how mosaicism is phenotypically manifested in the skeleton, we generated somatic mutants for *plod2* and *bmp1a*. In humans, mutations in *PLOD2* and *BMP1* are associated with Osteogenesis Imperfecta (OI), a heritable, heterogeneous group of connective tissue disorders commonly associated with skeletal deformities and bone fragility. OI is predominantly caused by dominant mutations in *COL1A1* or *COL1A2*, whereas mutations in non-collagenous genes mostly cause autosomal recessive forms of OI (18); the latter genes include *PLOD2* and *BMP1* (26). The enzyme encoded by *PLOD2*, lysyl hydroxylase 2, localizes to the endoplasmic reticulum, and catalyzes lysine residue hydroxylation in fibrillar collagen telopeptides (27). BMP1 is a secreted enzyme that functions in the cleavage of C-propeptides from procollagen precursors (28). We and others previously showed that zebrafish germline loss-of-function mutants for *plod2* and *bmp1a* exhibit severe skeletal abnormalities as adults, reminiscent of OI phenotypes (1, 27, 29, 30).

Somatic mutants for *plod2* and *bmp1a* were generated by injection of RNP complexes into embryos, and a subset of larvae were individually screened for indels at 12dpf. Sanger sequencing and TIDE analysis (31) revealed intra-animal mutation efficiencies of 82.7-88.1% and 71.0-87.5% for *plod2* and *bmp1a*, respectively. Adult phenotypes were visible at 90dpf, adding to several recent reports (11, 21) examining the durability of crispant zebrafish phenotypes through the larval-to-adult transition. At 90dpf, somatic mutants for both genes exhibited clear skeletal abnormalities similar to their adult germline mutant counterparts (Fig 3). Somatic mutants for *plod2* exhibited severe vertebral malformations including compression of the vertebrae along the anteroposterior axis, kyphosis, and increased bone formation. Somatic mutants for *bmp1a* exhibited increased vertebral radiopacity and bone thickening. Standard length (S.L.) was significantly reduced compared to sham controls for both *plod2* (control: 22.8mm [21.1-24.3mm], mutant: 19.8mm [17.0-22.6mm]; p=0.002; n=11/group; median [range]) and *bmp1a* (control: 23.8mm [21.8-24.4mm], mutant: 22.7mm [20.8-25.3mm]; p=0.009; n=15/group).

**Fig 3.**
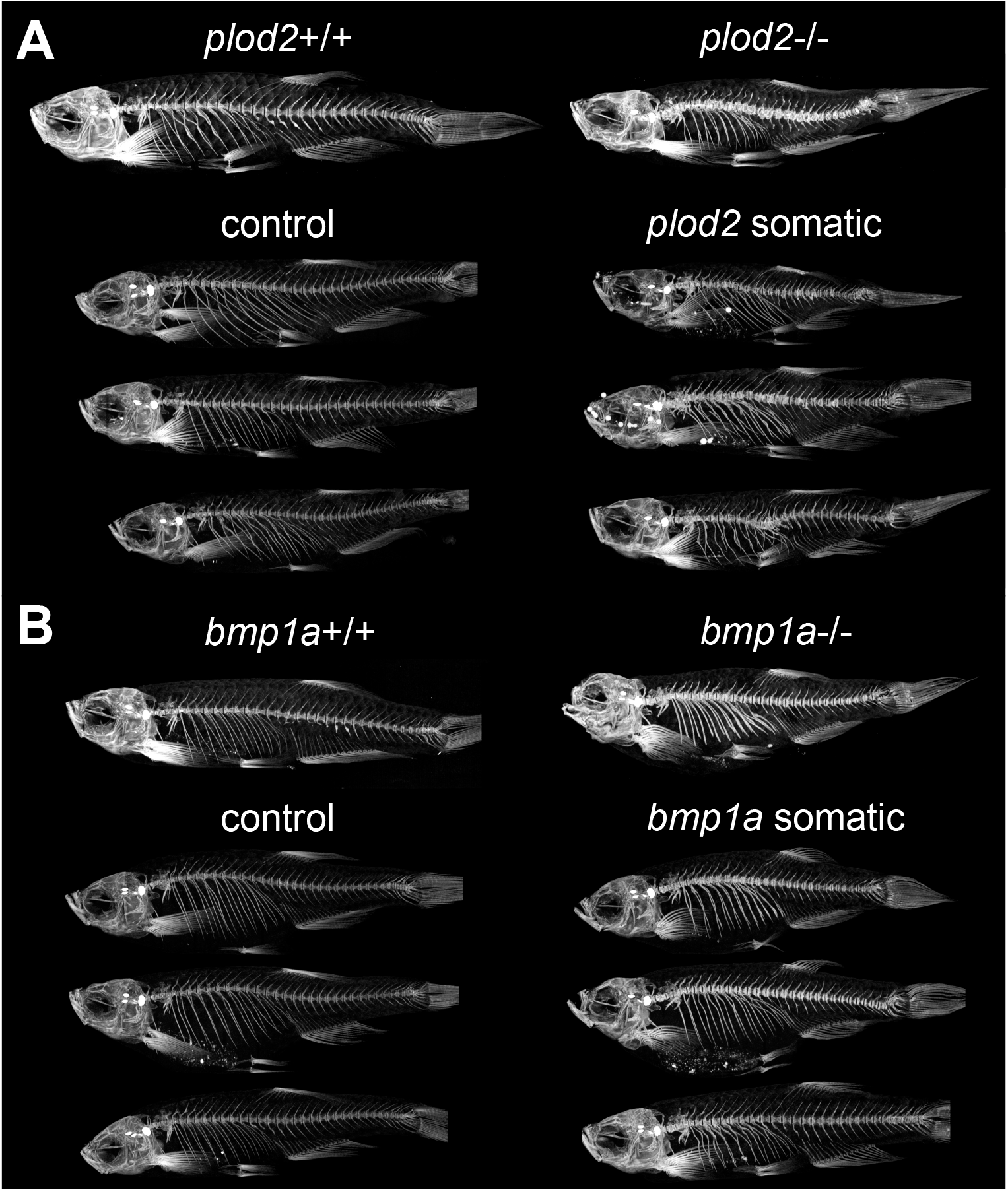
Somatic mutants for *bmp1a* and *plod2* exhibit inter-and intra-animal phenotypic variability. (A-B) Representative maximum intensity projections of microCT scans for *plod2* (A) and *bmp1a* (B) mutants.

Variability in phenotypic expressivity across animals was clearly evident. For *plod2*, such variability was perceptible by the number of dysmorphic vertebrae in each animal; in the 24 anterior-most precaudal and caudal vertebrae, *plod2* somatic mutants exhibited 12 [6-20] (median [range]) obviously thick or malformed vertebrae per fish. Further, 100% (11 out of 11) of animals were penetrant, as all individuals exhibited at least one severely malformed vertebra per animal. For *bmp1a* somatic mutants, most animals exhibited a qualitative increase in vertebral radiopacity and bone thickening. The extent of this increase was variable among individuals, ranging from mild to severe.

Variability in phenotypic expressivity within each animal was also evident in some cases. Certain *plod2* somatic mutants exhibited “patchy” expressivity characterized by contiguous spans of one or more dysmorphic vertebrae surrounded by vertebrae that appeared qualitatively normal. This pattern was reminiscent of spatial characteristics of macroscopic loss-of-function clusters in *sp7:EGFP* fish previously characterized. In *plod2* germline mutants, vertebrae are uniformly dysmorphic (27), suggesting that such “patchy” expressivity is not an inherent property of *plod2* loss of function in adult animals. In contrast to *plod2*, for *bmp1a* somatic mutants, intra-animal variability in phenotypic expressivity was less obvious; while radiopacity and thickening was variable from fish-to-fish, within each animal, these characteristics appeared to be relatively uniform. The relative uniformity in spatial expressivity in *bmp1a* crispants is discordant with the patchy mosaicism in osteoblasts of *sp7:EGFP* fish. Discordance between cell genotype and phenotype in mosaic animals suggets systemic/non-cell autonomous gene action. Consistent with this notion, the action of BMP1 is known to be non-cell autonomous, functioning as an extracellular enzyme involved in collagen processing. In contrast to the extracellular action of BMP1 in humans, the action of PLOD2 is intracellular as it modifies collagen in the endoplasmic reticulum.

### MicroCT-based phenomics enhances sensitivity in discriminating somatic mutant phenotypes

To characterize phenotypes quantitatively, we performed microCT-based phenomics (1). Recently, we developed a microCT-based workflow, FishCuT, which enables rapid (<5min/fish) quantification of 100s of measures in the axial skeleton of adult zebrafish (1, 18, 32). In this workflow, 25 different quantities are computed for each vertebra (Cent.TMD, Cent.Th, Cent.Vol, Cent.Le, Cent.SA, Cent.TMD.sd, Cent.Th.sd, Neur.TMD, Neur.Th, Neur.Vol, Neur.SA, Neur.TMD.sd, Neur.Th.sd, Haem.TMD, Haem.Th, Haem.Vol, Haem.SA, Haem.TMD.sd, Haem.Th.sd, Tot.TMD, Tot. Th, Tot.Vol, Tot.SA, Tot.TMD.sd, Tot.Th.sd; see (1) for description). Once calculated, these quantities are plotted as a function of vertebra number/level along the axial skeleton for each fish; we have termed such entities’vertebral traces’. For each combination of outcome/element, a standard score is computed as the difference between its value in each vertebral body and its mean value across all vertebrae in the control population, divided by the standard deviation across all vertebrae in the control population. These data are arranged into matrix constructs that we have termed ‘skeletal barcodes’. In our studies, 16 vertebrae (16*25=400 measures/animal) in 52 animals were analyzed, resulting in 52*400=20,800 data points that provided a comprehensive characterization of bone morphology and microarchitecture across the majority of the axial skeleton for each fish. To facilitate comparisons with prior studies (1), we present data on ten combinatorial quantities (the nine possible combinations of (Cent, HA, NA) × (Vol, TMD, and Th), plus Cent.Le) in the 16 anterior-most vertebrae in the main text, and have included all 25 quantities in the supplemental material.

Consensus methods to discriminate spatially varying phenotypes in somatic mutants have yet to be established. Previously, we showed that the global test (33), a regression-based statistical test designed for data sets in which many features have been measured for the same subjects, was effective in detecting differences in collections of vertebral traces in germline mutants (1). Statistical power in multivariate tests is dependent on underlying distributions—e.g., whether there are small changes in a large number of measures, or large changes in a few measures. In somatic mutants, only a subset of vertebra may be affected, and these vertebrae can be different from animal-to-animal. Thus, the performance of the global test in discriminating somatic mutant populations, and how it compares to univariate approaches, is unknown. Thus, we tested the hypothesis that assessing vertebral patterns with the global test would provide greater sensitivity in distinguishing somatic mutants with variable phenotypic expressivity compared to (a) Mann-Whitney (M.-W.) of individual vertebrae, and (b) M.-W. tests of quantities averaged across all vertebrae. We chose the M.-W. test as a reference univariate test because, like the global test, the M.-W. test is non-parametric.

To test this, we performed Monte Carlo simulations (See SI Appendix). The universal scaling distribution defined in *sp7:EGFP* somatic mutants (Eq 1) was used to simulate different patterns of mosaicism and levels of phenotypic variability (1,000 simulations per analysis). For microscopic loss-of-function clusters (Fig 4A), analyzing vertebrae 1:16 using the global test resulted in up to a 1.41-fold increase in sensitivity (fraction of times in which p<0.05 when comparing simulated mutant fish to WT fish) compared to using the M.-W. test using vertebra 2, and a 1.15-fold increase compared to the M.-W. test using quantities averaged across vertebrae 1:16 (Fig 4B). Differences in testing procedure were dependent on how loss-of-function regions were spatially clustered. For instance, for macroscale loss-of-function clusters (Fig 4C), analyzing Cent.Vol in vertebrae 1:16 with the global test conferred up to a 3.65- and 1.21-fold increase in sensitivity compared to analyzing vertebra 2 and the mean of vertebrae 1:16 with the M.-W. test, respectively; noticeably higher compared to simulations using microscale clusters (Fig 4D).

**Fig 4.**
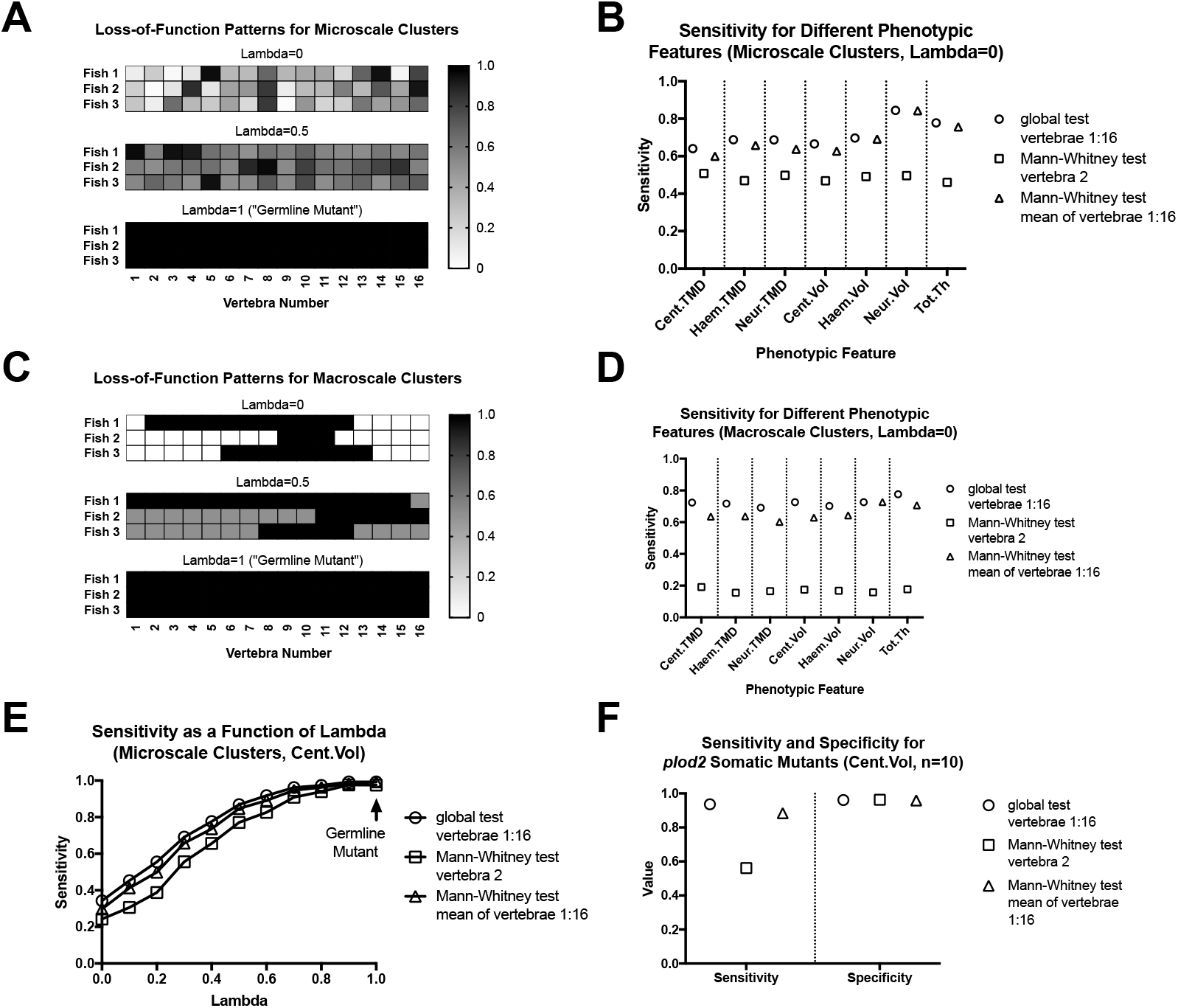
Analyzing vertebral patterns using the global test increases sensitivity in discriminating somatic mutants compared to analyzing single readouts. (A) Representative mosaic patterns for microscale loss-of-function clusters. Values indicate extent of gene loss. (B) Sensitivity of the global test compared to univariate approaches for somatic mutants with microscale loss-of-function clusters. (C) Representative mosaic patterns for macroscale loss-of-function clusters. Values indicate extent of gene loss. (D) Sensitivity of the global test compared to univariate approaches for somatic mutants with microscale loss-of-function clusters. (E) Sensitivity of the global test in detecting changes in centrum volume compared to univariate approaches for somatic mutants with microscale loss-of-function clusters with different degrees of intra-animal variability (parameterized by lambda). Relative differences between the global test and univariate tests become heightened as mosaicism increases (i.e., as lambda goes to zero). (F) Results for bootstrap simulations using phenotypic profiles from *plod2* somatic mutants.

Notably, we found that the relative benefits of the global test, compared to the univariate tests, became heightened as mutants become increasingly mosaic (i.e., less germline mutant-like). Specifically, the sensitivity of the global test increased relative to the other tests with decreased values of lambda, the model parameter which parameterizes intra-animal variation (Fig 4E). While we did not explicitly vary characteristic effect size, we previously showed that the relative benefits of the global test, compared to t-tests, are highest as smallest effect size (1). Finally, we performed non-parametric simulations using experimental data derived from *plod2* somatic mutants (Fig 4F). Consistent with our analyses in parameterized simulations, the global test conferred higher sensitivity, with similar specificity, compared to M.-W. tests using averaged quantities as well as individual vertebrae. For all simulations, specificity (1 - the fraction of times in which p<0.05 when comparing WT to WT fish) ranged between 0.94-0.97, closely bracketing the expected value of 0.95. Taken together, our studies show that the global test is an effective test for detecting differences in collections of spatially varying phenotypes in somatic mutants. Further, they provide evidence that deep phenotyping increases sensitivity, with similar specificity, in discriminating somatic mutant populations, compared to analyzing single readouts.

### MicroCT-based phenomics can discriminate differences in spatial phenotypic variability

Spatial phenotypic variation in mosaic individuals has the potential to encode biological information. This is demonstrated by the high and low spatial variability seen in *plod2* and *bmp1a* somatic mutants, respectively, which we speculate is due to differences in autonomy of gene action. Yet, consensus approaches to detect differences in spatial phenotypic variability are lacking. We explored the utility of Moran’s I, a measure of global spatial autocorrelation commonly employed for geostatistical analysis, for this purpose. Moran’s I usually ranges from approximately −1 to 1, and can be interpreted as the extent to which values are spatially clustered (positive), dispersed (negative), or random (zero) (Fig 5A). In Monte Carlo simulations, we found that microscale clusters resulted in Moran’s I tending to decrease, whereas macroscale clusters resulted in Moran’s I tending to increase (Fig 5B). For *plod2* and *bmp1a* somatic mutants, when the distribution of Moran’s I was calculated across all 10 combinatorial measures, there was a marked shift in the center of the distribution toward I=0 for *plod2* somatic mutants compared to controls (Fig 5C). In contrast, no obvious shift in distribution center was observed for Moran’s I in somatic mutants for *bmp1a* (Fig 5D), consistent with our qualitative observations. Thus, we conclude that Moran’s I is an effective measure of spatial phenotypic variation, which may be useful for situations where qualitative observation of spatial variation is not possible. Moreover, because shifts in Moran’s I in our simulations could be observed in the presence of relatively small effect sizes (e.g., 2-3× smaller than those in our study), our simulations suggest that Moran’s I may be a useful metric to help discriminate differences in spatial variability for mutants with more subtle phenotypic effects. In Fig 5–Supplemental Fig 1, we provide further analyses explaining the numerical basis of changes in Moran’s I observed in our simulations.

**Fig 5.**
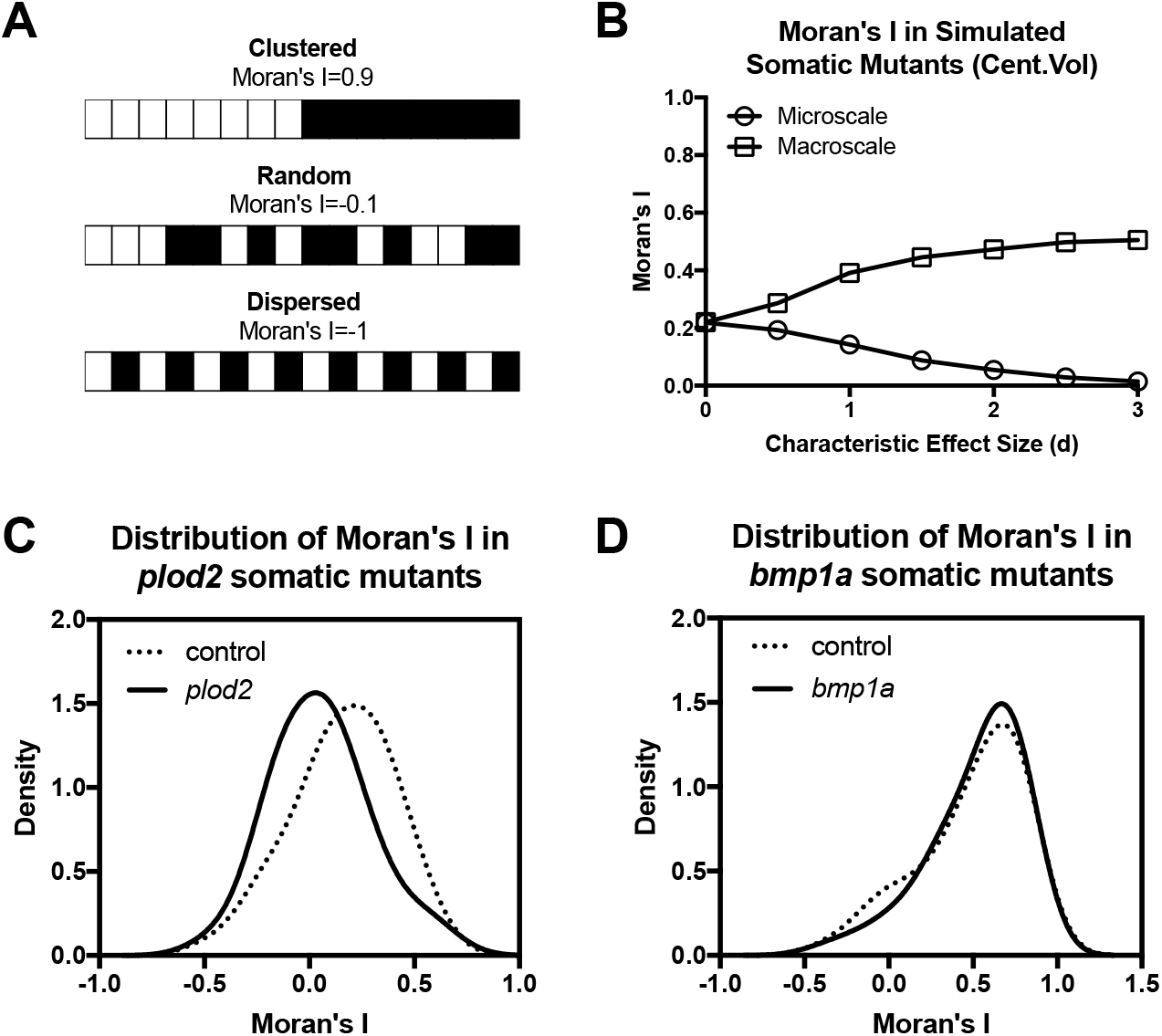
Quantification of spatial phenotypic variation in somatic mutants. (A) Schematic demonstrating Moran’s I for different patterns of mosaicism. (B) Changes in Moran’s I with characteristic effect size, d. Microscale loss-of-function clusters decrease Moran’s I, whereas macroscale clusters increase it. (C,D) Moran’s I for *plod2* mutants (C) shows a shift towards a random distribution compared to controls, while that of *bmp1a* mutants (D) does not.

### Somatic and germline mutant phenomes for plod2 and bmp1a exhibit high correspondence

We returned our attention to *plod2* and *bmp1a* somatic mutants, and assessed which FishCuT measures exhibited differences in the global test. Analysis for *plod2* somatic mutants (n=11 fish/group) exhibited significant differences in centrum, haemal arch, and neural arch tissue mineral density (Cent.TMD: p=0.000005, Haem.TMD: p=0.00008, Neur.TMD: p=0.00005); centrum volume (p=0.004), thickness (p=0.04), and length (p=0.00009); and neural arch thickness (p=0.00007) (Fig 6). Somatic mutants for *bmp1a* (n=15 fish/group) exhibited significant differences in centrum, haemal arch, and neural arch tissue mineral density (Cent.TMD: p=0.000004, Haem.TMD: p=0.00002, Neur.TMD: p=0.00005); centrum volume (p=0.004), thickness (p=0.02), and length (p=0.01); and haemal arch thickness (p=0.008) (Fig 7). Data for all 25 combinatorial measures for *plod2* and *bmp1a* are provided in Fig 6–Supplemental Fig 1 and Fig 7–Supplemental Fig 1, respectively. For both *plod2* and *bmp1a*, most significantly different features were associated with vertebral traces that were elevated or depressed across all vertebrae; an exception was neural arch thickness in *plod2* somatic mutants, which was lower in anterior vertebrae, but higher in posterior vertebrae.

**Fig 6.**
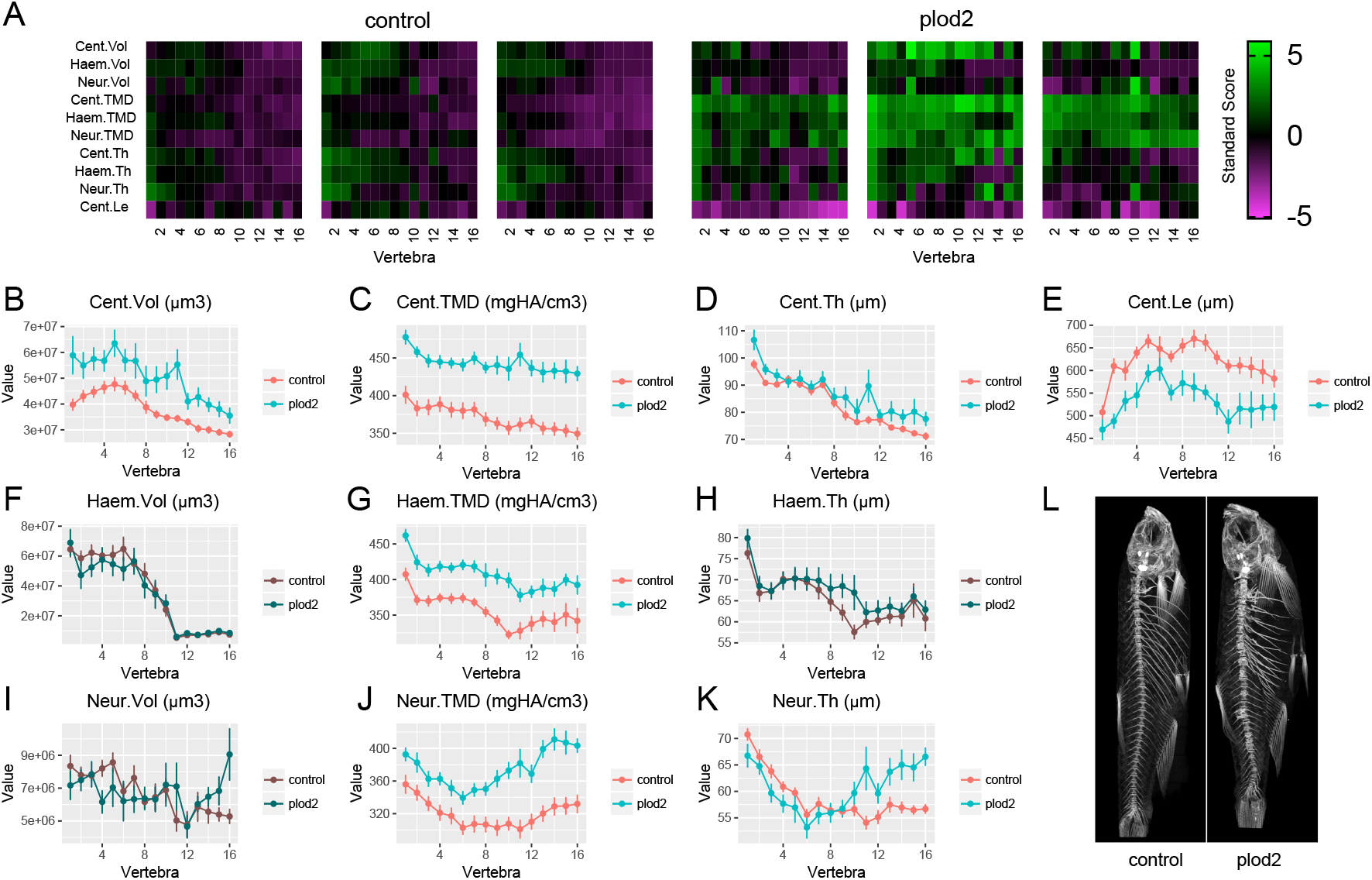
FishCuT analysis of *plod2* somatic mutants. (A) Skeletal barcodes for control and *plod2* somatic mutant fish. Each barcode represents a single fish (n = 3 fish per group shown). Standard scores are computed as the difference between the value of the feature in the individual and the mean value of the feature across all vertebrae in the control population, divided by the standard deviation of the feature across all vertebrae in the control population. (B–K) Phenotypic features plotted as a function of vertebra (mean ± SE, n = 11/group). Plots associated with p<0.05 in the global test are colored in a lighter coloring scheme (see text for p-values). (L) Maximum intensity projections of microCT scans.

**Fig 7.**
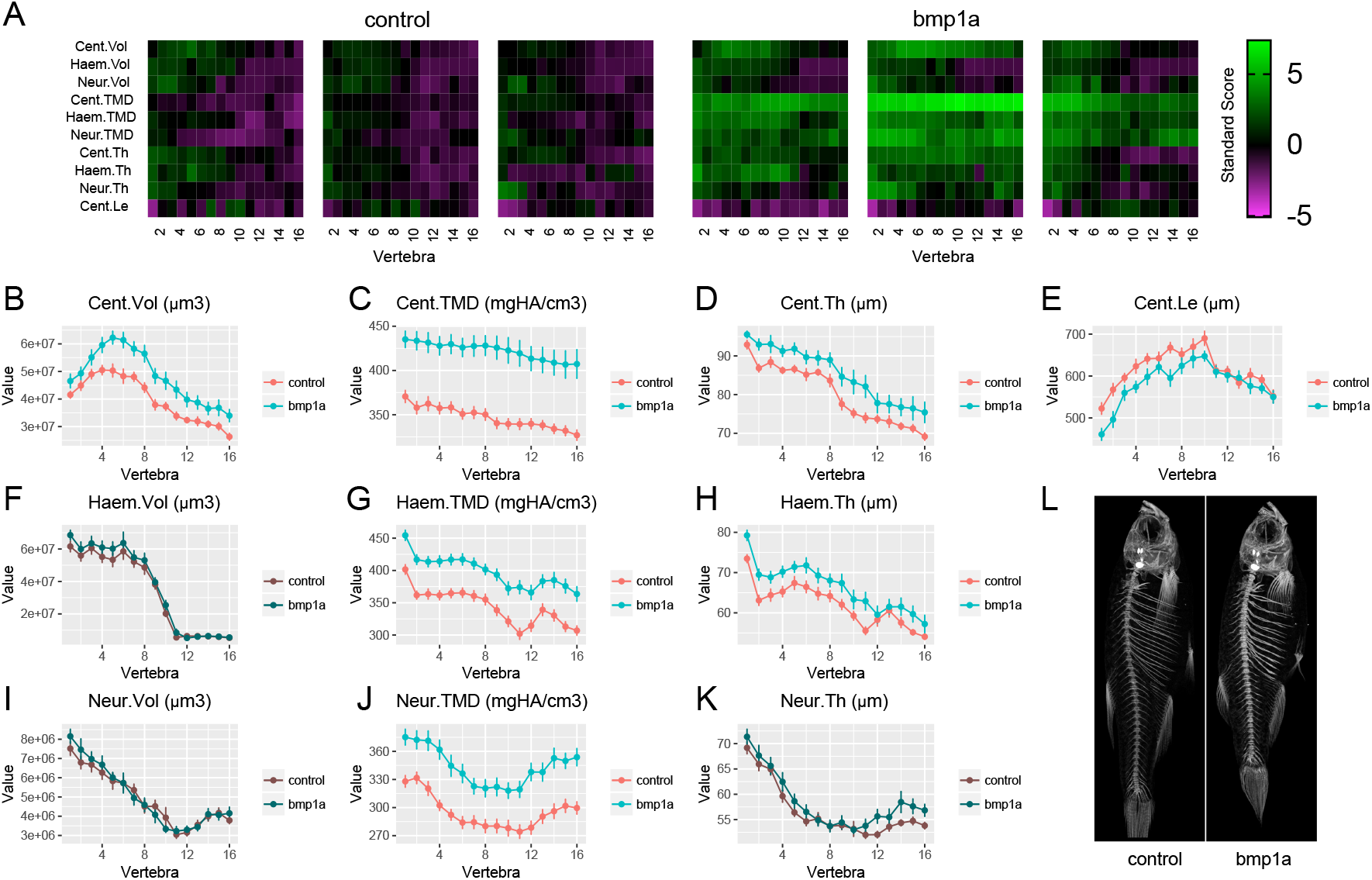
FishCuT analysis of *bmp1a* somatic mutants. (A) Skeletal barcodes for control and *bmp1a* somatic mutant fish. Each barcode represents a single fish (n = 3 fish per group shown). Standard scores are computed as the difference between the value of the feature in the individual and the mean value of the feature across all vertebrae in the control population, divided by the standard deviation of the feature across all vertebrae in the control population. (B–K) Phenotypic features plotted as a function of vertebra (mean ± SE, n = 15/group). Plots associated with p<0.05 in the global test are colored in a lighter coloring scheme (see text for p-values). (L) Maximum intensity projections of microCT scans.

Next, we assessed the extent to which somatic mutants for *plod2* and *bmp1a* could act as faithful models of their germline mutant counterparts. For these analyses, we compared global test results to measurements and FishCuT outputs for *plod2* and *bmp1a* germline mutants (n=3 for both germline groups) previously generated in (1) (Fig 8). Characteristic effect sizes in *plod2* and *bmp1a* somatic mutants were, on average, 25.6% and 24.4%, respectively, of those in their germline mutant counterparts (*plod2* somatic: 0.95; *plod2* germline: 3.71; *bmp1a* somatic: 0.80; *bmp1a* germline: 3.23). Occurrence of outliers (defined as absolute barcode values residing >1.5× the interquartile range (IQR) from the median (34)) were not appreciably different across somatic and germline mutant groups (*plod2* somatic: 1.0%, *plod2* germline: 7.3%; *bmp1a* somatic: 1.1%, *bmp1a* germline: 0.6%).

**Fig 8.**
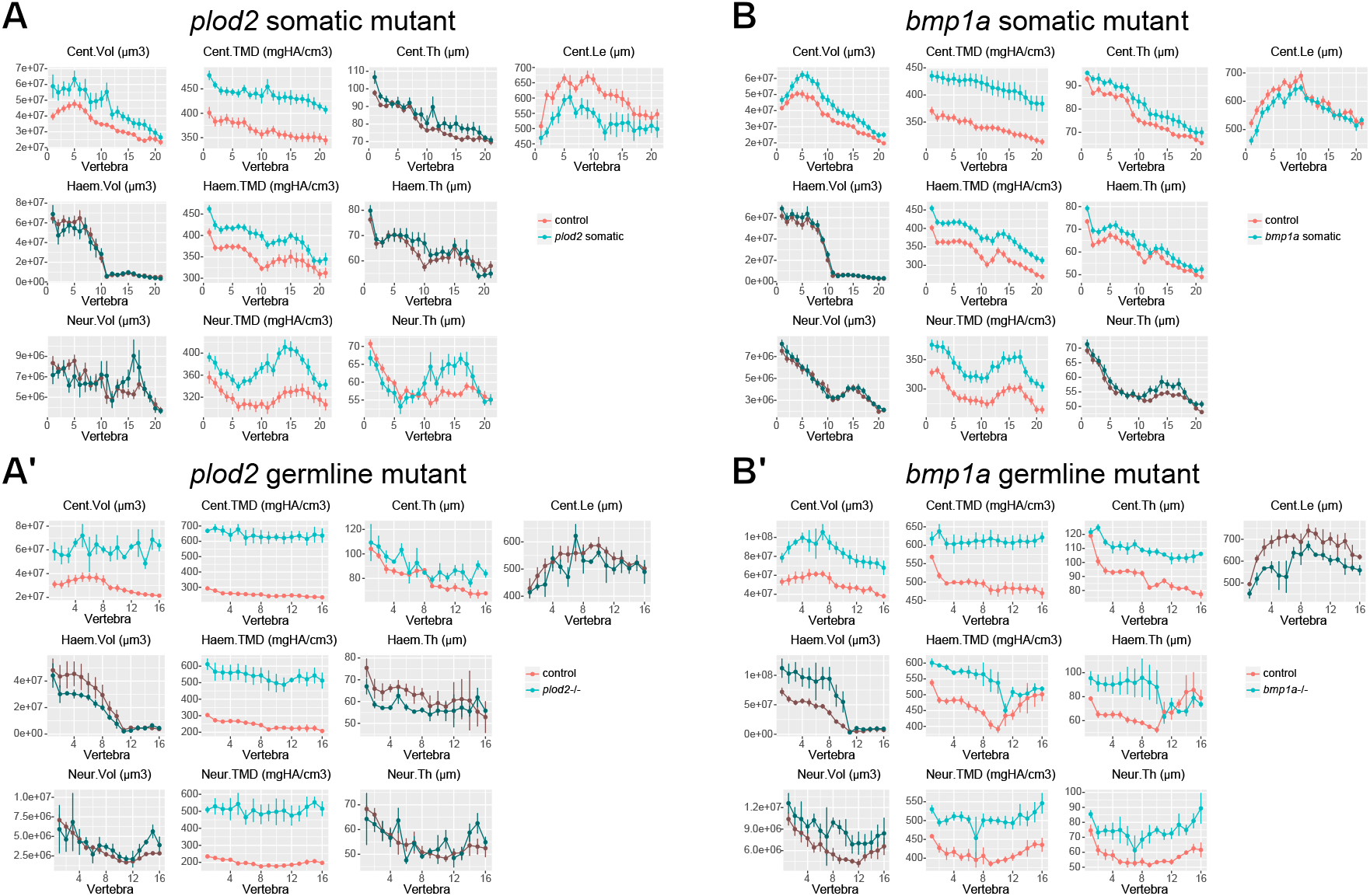
Correspondence in between somatic and germline mutant phenotypes for *plod2* and *bmp1a*. Phenotypic features are plotted as a function of vertebra (mean ± SE). Plots associated with p<0.05 in the global test are colored in a lighter coloring scheme (see text for p-values). (A-A’) Comparison of *plod2* somatic (A) and germline (A’) mutants. Data in (A’) were subjected to allometric normalization, due to the severe reduction in body length in *plod2* germline mutants. (B-B’) Comparison of *bmp1a* somatic (B) and germline (B’) mutants. High correspondence in significantly different measures is observed between somatic and germline mutants for both genes. For A’ and B’, n=3/group.

Since the reduced standard length in *plod2* somatic mutants was more muted compared to *plod2* germline mutants (~4× less), we compared somatic mutant results to *plod2* germline mutant phenotypes that had been subjected to allometric normalization. We previously showed that by transforming WT sibling data to a ‘virtual’ phenome scaled to the mean standard length of age-matched mutants, allometric models provided a means to enable length-matched fish comparisons from an age-matched control group (1). Across the 10 primary measures, somatic mutants for *plod2* exhibited significant differences for 80% (4 out of 5) of the measures significantly altered in *plod2* germline mutants (Fig 8A,A’). Moreover, 60% (3 out of 5) of the combinatorial measures not significantly different in *plod2* mutants were also not different in *plod2* somatic mutants. Correspondence was noticeably lower when *plod2* somatic mutants were compared to *plod2* germline mutants that had not been allometrically normalized (3 of 6 corresponding measures with statistical significance, 50%; and 1 of 4 corresponding measures without statistical significance, 25%). Comparisons of results to the other 15 measures in FishCuT were not possible because allometric models have yet to be developed for them.

We also observed a high degree of correspondence between *bmp1a* somatic and germline mutant phenotypes (Fig 8B,B’). Specifically, across the 10 primary measures, *bmp1a* somatic mutants exhibited significant differences for 86% (6 out of 7) of the measures significantly different in *bmp1a* germline mutants. Further, 67% (2 out of 3) of the measures not significantly different in *bmp1a* germline mutants were also not significantly different in *bmp1a* somatic mutants. Correspondence was not improved when comparing *bmp1a* somatic mutants to *bmp1a* germline mutants that had been subjected to allometric normalization. These studies suggest the potential for somatic mutants to predict germline mutant phenotypes with high fidelity.

Notably, for *plod2* somatic mutants, mean vertebral traces appeared “smooth”— despite intra-individual variability in phenotypic expressivity in some measures (Fig 8-Supplemental Fig 1)—due to averaging across the sample. Moreover, the magnitude of standard errors in somatic mutants, relative to differences in the mean, approached or were reduced compared to those in germline mutants, due to the larger sample sizes in somatic mutant groups (*plod2* somatic: n=11/group, *plod2* germline: n=3/group; *bmp1a* somatic: n=15/group, *bmp1a* germline: n=3/group). In our Monte Carlo simulations, somatic mutants with characteristic effect sizes similar to those in *plod2* and *bmp1a* somatic mutants were detected in the global test with >80% power (assuming alpha=0.05) using the sample sizes in our study. Taken together, somatic mutants can exhibit high correspondence with germline mutants, provided a sufficiently large sample, the size of which can be rationally estimated from simulations.

Somatic mutant analysis identifies skeletal phenotype following loss-of-function in wnt16 While the above studies suggested the benefits of deep skeletal phenotyping in increasing the fidelity of CRISPR-based G0 screens, one limitation is that they were performed in zebrafish models of monogenic bone disorders. We investigated consequences of loss-of-function in *wnt16*, a gene which has strong ties to genetic risk for osteoporosis in humans (35). In mice, loss of WNT16 affects both osteoclastogenesis (35) and bone formation (36, 37). WNT16 knockout in mice can result in low incidence of spontaneous fractures, depending on genetic background (38). However, in most mutants, bone appears grossly normal, with no trabecular loss, and moderate cortical thinning that can be resolved by microCT. Thus, we generated *wnt16* somatic mutants in zebrafish to establish proof-of-concept that, by enhancing sensitivity in discerning somatic mutant phenotypes, G0 phenomic profiling may facilitate the detection of genes that are relevant to osteoporosis, a disease whose complex, multigenic nature has hampered the identification of causal variants and gene targets.

Somatic *wnt16* mutants were generated in the same manner as those for *bmp1a* and *plod2*. TIDE analysis revealed that *wnt16* mutation efficiencies for individual 12dpf larvae ranged from 67.4% to 93.6% with a median efficiency of 80.1%. Mutational efficiency at this site was confirmed using next generation sequencing (NGS) (Fig 9-Supplemental Fig 1). At 90dpf, microCT scans revealed variable phenotypic penetrance and expressivity (Fig 9). Standard length was significantly reduced in *wnt16* somatic mutants compared to controls (control: 22.0mm [17.6-25.1mm], mutant: 18.8mm [16.0-22.6mm]; p=0.02; n=12/group; median [range]).

**Fig 9.**
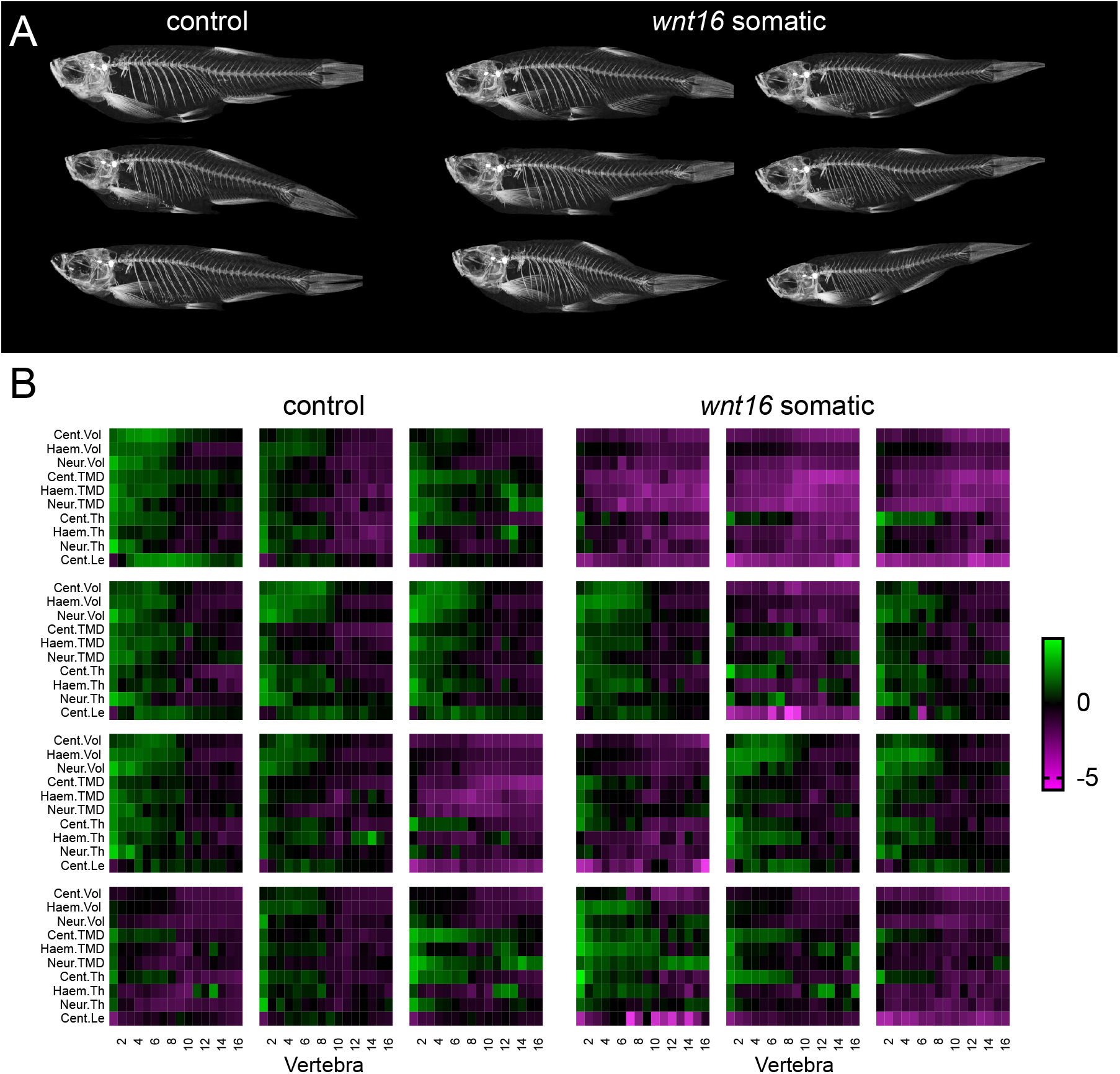
Somatic *wnt16* mutants show variable expressivity. (A) Representative MicroCT maximum intensity projections of *wnt16* somatic mutants. (B) Skeletal barcodes for control and *wnt16* somatic mutant fish. Each barcode represents a single fish (n = 12 fish per group shown). Standard scores are computed as the difference between the value of the feature in the individual and the mean value of the feature across all vertebrae in the control population, divided by the standard deviation of the feature across all vertebrae in the control population.

Somatic mutant phenotypes for *wnt16* were more subtle compared to those for *plod2* and *bmp1a*. Characteristic effect sizes were reduced for somatic mutants for *wnt16* compared to the other two genes (*wnt16* somatic: 0.72; *plod2* somatic: 0.95; *bmp1a* somatic: 0.80). Moreover, *wnt16* somatic mutants exhibited variability in body size and gross appearance that made robust phenotypic changes—specifically, those that were clearly independent of differences in developmental progress—difficult to discern by eye.

FishCuT analysis revealed a low bone mass phenotype in *wnt16* somatic mutants. Somatic mutants for *wnt16* exhibited significant changes in centrum volume (p=0.02) and centrum length (p=0.004) compared to sham injected clutchmates (n=12/group; Fig 10A). Following allometric normalization, significant differences were observed for centrum TMD (p=0.0001), haemal arch TMD (p=0.0001), neural arch TMD (p=0.0002), and neural arch thickness (p=0.03) (Fig 10-Supplemental Fig 1). Because *wnt16* somatic mutants did not exhibit normal allometry with standard length, this suggests that morphological differences in *wnt16* somatic mutants were not solely due to developmental delay. Evaluation of morphological defects (rib fracture calluses, neural arch non-unions, centrum compressions, and centrum fusions) revealed no significant differences in the occurrence of a single defect type in *wnt16* somatic mutants compared to controls. However, the proportion of animals exhibiting any such defect was significantly higher in *wnt16* somatic mutants (p=0.01, n=12/group).

**Fig 10.**
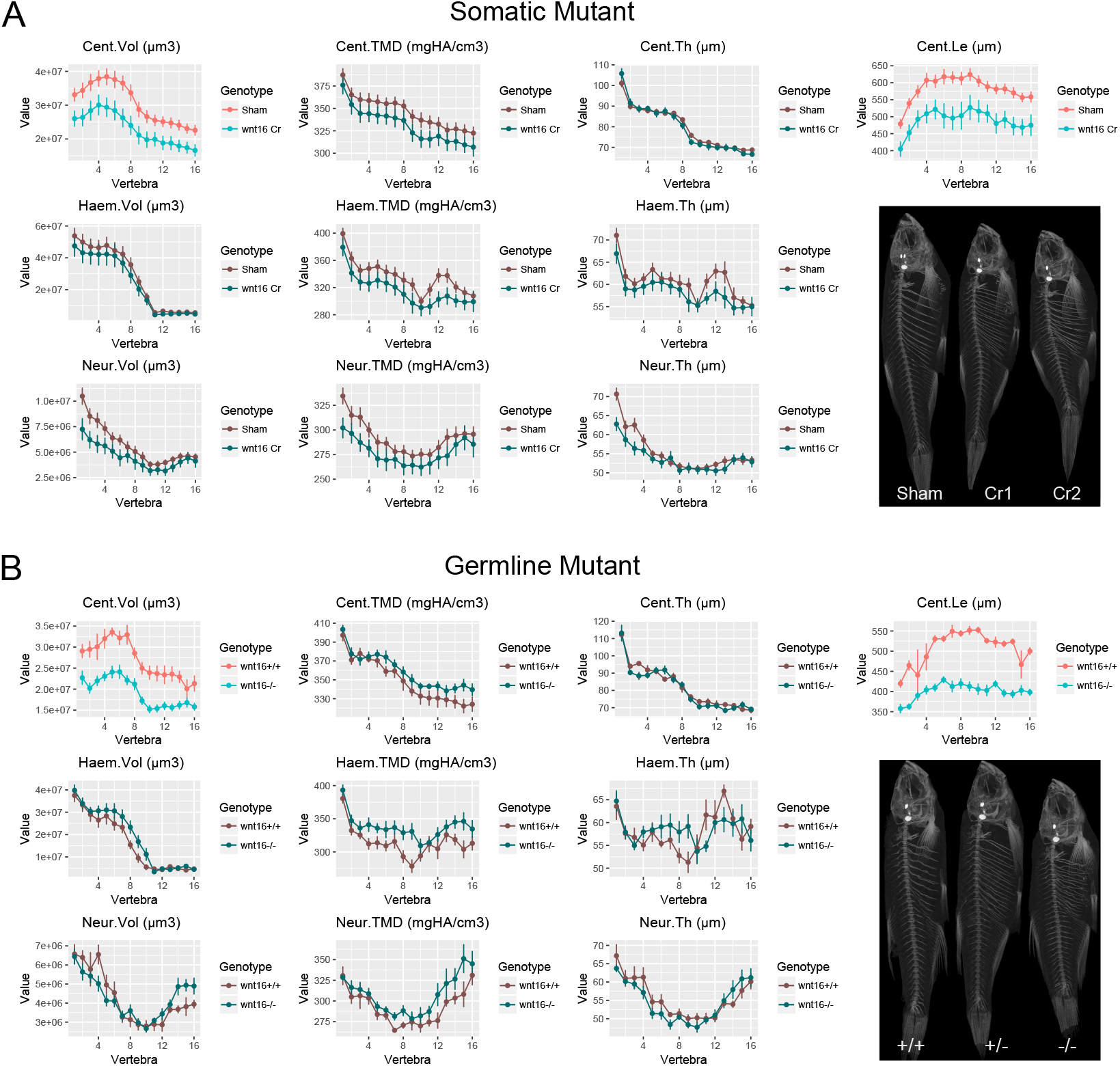
FishCuT analyses of *wnt16* somatic and germline mutants. In Figures A-B, phenotypic features are plotted as a function of vertebra (mean ± SE). Plots associated with p<0.05 in the global test are colored in a lighter coloring scheme (see text for p-values). (A) Analysis of *wnt16* somatic mutants. Somatic *wnt16* mutants show decreased centrum volume and centrum length compared to sham injected controls (n=12/group). Maximum intensity projections of microCT scans of representative sham fish (sham) and two somatic mutants displaying different expressivity (Cr1, Cr2) are shown. (B) Analysis of *wnt16* germline mutants. Germline *wnt16*-/- mutants show decreased centrum volume and centrum length compared to *wnt16*+/+ controls (+/+: n=4, -/-: n=6). Maximum intensity projections of microCT scans of representative fish of each genotype are shown.

Wnts are secreted molecule that have been shown to exhibit long-range diffusion (39). Because of our studies in *bmp1a* somatic mutants suggesting that mutations in genes encoding for secreted factors result in lower spatial variability relative to those that whose actions are cell autonomous, we predicted that spatial phenotypic variability in *wnt16* somatic mutants would be relatively low, and comparable with WT animals. Consistent with this notion, when we examined the distribution of Moran’s I in *wnt16* somatic mutants, no obvious difference was observed compared to controls (Fig 10-Supplemental Fig 2).

Finally, to determine if the somatic phenotype is representative of germline dysfunction in *wnt16*, we generated *wnt16* F2 germline mutants harboring a frameshift allele (a 10 base pair insertion in exon 3) and subjected these fish to phenomic profiling (n=4 +/+, n=6 +/−, and n=6 -/-). Germline *wnt16*-/-mutants exhibited significant changes in centrum volume (p=0.002) and centrum length (p=7.1e-6) compared to +/+ clutchmates (Fig 10B). Heterozygous *wnt16* germline mutants showed no significant changes compared to controls (Fig 10-Supplemental Fig 3), suggesting that the somatic *wnt16* phenotype arises from mosaic bialleleic loss-of-function. Standard length was also significantly reduced in homozygous mutants (+/+: 17.4mm [15.8-17.9mm], -/-: 20.3mm [18.9-20.8mm]; p=0.01; median [range]). No obvious differences in the frequency of centrum compressions and neural arch non-unions were observed between groups (although, germline mutant studies were underpowered to detect morphological defects). Notably, the phenotype seen for *wnt16* mutants was concentrated in the centrum (somatic and germline). In teleosts, centrum growth through the larval-to-adult transition has been attributed to osteoblasts originating in the intervertebral region (40).

These somatic *wnt16* data, together with the germline correspondence for *plod2* and *bmp1a* somatic mutants, support the fidelity of phenotyping directly in CRISPR-edited G0 zebrafish for increased throughput in genetic screens, despite variability arising from CRISPR-induced mutations and the potential for unintended off-target effects. Further, our results identify impaired bone mass accrual in adult zebrafish arising from both somatic and constitutional loss-of-function in *wnt16*, a gene implicated in heritable risk for osteoporosis, adding to evidence that there are contexts in which osteoporosis-related traits can be identified and modeled in zebrafish for the study of genes implicated as risk factors in this disease.

## DISCUSSION

Next generation phenotyping technologies hold promise to open new opportunities to understand and exploit somatic mutant biology. In this study, we described evidence of 1) of a universal distribution describing sizes of loss-of-function clusters arising from CRISPR editing, 2) different spatial patterns caused by somatic mutations in genes associated with cell-autonomous and non-cell autonomous actions, and 3) effective statistical methods for analyzing spatially variable phenotypes. By advancing our understanding how somatic mutations disseminate during development, how these mutations manifest as phenotypic patterns, and how to quantitatively detect differences in these patterns within phenomic datasets, our studies help in decoding somatic mutant phenotypes using next generation phenotyping.

By visually depicting functional loss in *sp7*+ cells, we identified distinct characteristics in regard to how loss-of-function cell clusters in the skeleton distribute in space and size following CRISPR editing. We found that Eq 1, which was derived based on modeling of fluorescence-based lineage tracing studies, described loss-of-function cluster sizes in skeletal tissues derived from both the sclerotome (vertebrae), and neural crest (branchiostegal rays), and in animals with different levels of mutational loss-of-function. Because this relation arises via contributions from clonal fragmentation and merger events rather than cell fates specified by developmental programs, the data collapse in our study is unlikely to be unique to skeletal tissues, as clonal fragmentation and merger events initiate in the early developing embryo (24). While it remains to be seen whether a similar size distribution in loss-of-function clusters is observed in other tissues and organs, in lineage tracing studies, Eq 1 has been found to fit clonal cluster size distributions in diverse contexts including during development of the mouse and zebrafish heart, the mouse liver, and the mouse pancreas (24). Notably, this size distribution is not observed in developmental events where clonal fragmentation and merger events are suppressed (e.g., mouse acinar cells (24)), and we would expect similar conditionality to apply in regard to CRISPR-induced somatic mutations. Still, a general relationship describing loss-of-function cluster sizes in somatic animals may facilitate the interpretation of spatially varying phenotypes commonly observed in G0 screens. This may also have value in other instances where estimates of cluster size distributions may be needed, such as in therapeutic applications of CRISPR editing.

While our studies implicate clonal fragmentation and merger as an influential factor in mosaic pattern development, they do not imply that developmental programs are unimportant. Indeed, within individual fish, we observed spatial patterns that were clearly non-random, and which could be explained by models of teleost spine development. In zebrafish, the centrum first ossifies through direct mineralization of the notochord sheath, which is then encased by intramembranous bone produced by somite-derived osteoblasts (41). In medaka, *twist*+ sclerotomal cells concentrate in the intervertebral regions prior to centrum ossification, and supply osteoblasts to the centra flanking either side of the intervertebral region, as well as the adjacent neural arches (40). In this intervertebral growth center (IGC) model, osteoblasts on adjacent centra, as well as the arches they flank, are all descendants of the same intervertebral cells. The IGC model predicts that loss-of-fluorescence should be related in the centra of adjacent vertebra, as well as in the neural arches and centra of the same vertebral bodies—events we observed in individuals. In this context, developmental programs and clonal fragmentation and merger events may work in concert to influence mosaic patterns; while only the latter contributes to size distributions in loss-of-function clusters, both contribute to spatial distributions. Finally, because bone tissue accumulates throughout life, it is possible that spatial variations in somatic mutant skeletal phenotypes arise from evolving spatial patterns in mosaicism during growth. This can potentially be tested by visualizing patterns of CRISPR-induced lesions longitudinally. However, *in vivo* fluorescent imaging of the spine is difficult in adults, even in mutants that lack pigmentation, due to the scattering of light in deep tissue.

As a spatially distributed system comprised of repeating anatomical units, the spine is an excellent substrate for understanding how mosaicism manifests as spatial phenotypic patterns. We found that mutant phenotypes could be expressed relatively uniformly in the spine (*bmp1a, wnt16*), or in alternating patterns of affected and unaffected vertebrae (*plod2*). The relatively uniform phenotypic expressivity in *bmp1a and wnt16* somatic mutants was discordant with the patchy loss-of-fluorescence patterns observed in *sp7:EGFP* fish. Indeed, phenotypic variability resembling a dosage curve has been previously hypothesized to occur when mutating a gene encoding a secreted factor, with different numbers of cells carrying the relevant mutation (42). Our studies suggest that somatic mutants can yield phenotypes for genes associated with cell autonomous or non-cell autonomous function, though the former may be associated with higher spatial variation. One practical implication is that for germline mutants caused by genes with unknown function, somatic mutants may be intentionally created, and screened for spatial phenotypic variability, as a means to inform autonomy in gene action.

Statistical methods to decode somatic mutant phenotypes are lacking. We found that the global test was an effective test in discriminating collections of spatially variable phenotypes. The increased sensitivity of analyzing vertebral patterns with the global test, relative to univariate approaches, underscore the potential for deep phenotyping to improve G0 screen productivity, and facilitate the study of genetic variants of smaller effect sizes. Notably, the global test can be used for multivariate phenotypes where spatial relationships are not specified (e.g., a panel of measures taken at the same anatomical site), and thus may be broadly useful for detecting differences in groups of measures between different populations. We also found that Moran’s I was sensitive to spatial phenotypic variation, suggesting that it could be employed in other G0 screens to extract additional biological information, and should be generalizable to other organ systems and tissue types. For example, in a previous G0 screen, Shah et al. enumerated the presence of 30 electrical synapses along the spinal cord, and used these data to calculate % loss of synapses in each animal (10). In principle, the spatial distribution of abnormal synapses could be analyzed using Moran’s I, to inform cell autonomy in gene action. Alternatively, clusters of abnormal synapse could be identified, their size distributions enumerated, and compared to Eq 1 to assess conformance to the expected distribution of loss-of-function mutations.

Technologies mirroring those in our studies may be useful for clinical diagnosis. For example, in human patients with mutations in COL1A1, a burden of more than 40%- 75% of osteoblasts containing at least one mutant allele becomes incompatible with normal skeletal function (43). Whether mosaicism in different forms of OI tends to manifest as “patchy” or relatively uniform phenotypic expressivity is unknown, as intra-individual variability in phenotypic expressivity in OI is, in general, not rigorously examined. However, there are instances of localized skeletal deficiencies, such as proximal femoral focal deficiency (PFFD), in which ~85-90% of cases are unilateral (44). Familial cases for PFFD have been described (45). By showing differences in Moran’s I between *plod2* somatic mutants and controls, our studies suggest the potential for deep phenotyping, in concert with spatial statistical analysis, to help distinguish phenotypic variation that naturally occurs within individuals, from that associated with mosaicism.

Human genome-wide association studies have identified over 200 loci (46, 47) associated with BMD and fracture risk in humans, however, most of the causal genes responsible for these associations have yet to be definitively assigned. Large-scale screens of candidate genes provide a mechanism to attribute functional skeletal contributions of these genes to osteoporosis-related traits. We identified a bone mass phenotype in zebrafish somatic and germline mutants for *wnt16*. Because this gene is implicated in mediating inter-individual variation in BMD and fracture risk in humans (48, 49), this broadens our understanding of zebrafish mutant phenotypes that may be predictive of human BMD and fracture risk. Somatic *wnt16* mutants successfully recapitulated phenotypic signatures in germline *wnt16* mutants, providing further evidence of the fidelity of phenotyping directly in CRISPR-edited G0 somatic mutants. While throughput of our workflow is limited by the time required for microCT acquisition (~5min/fish when 8 fish are scanned simultaneously (1)), this can potentially be overcome advances in high-throughput microCT (1). Though it remains to be seen if faithful correlation between somatic and germline mutant phenotypes exists for most skeletal genes, microCT-based phenomics, in concert with CRISPR-based G0 screens, is a promising direction for the rapid identification of candidate genes and development of targeted therapies for mono- and multigenic skeletal diseases.

In conclusion, our studies advance our understanding of how somatic mutations disseminate during development, how these mutations manifest as phenotypic patterns, and how to quantitatively detect differences in these patterns within phenomic datasets. In doing so, our studies help in using next generation phenotyping to decode somatic mutant phenotypes, which may confer advantages and additional insights in both experimental and clinical contexts.

## MATERIALS AND METHODS

For more detailed methods, see SI Appendix.

### Zebrafish rearing

All studies were performed on an approved protocol in accordance with the University of Washington Institutional Animal Care and Use Committee (IACUC). Studies were conducted in mixed sex animals from either the wild-type AB strain or the transgenic *sp7:EGFP* (23) background. A description of *plod2* and *bmp1a* germline strains used for comparative analysis is described in (1).

### CRISPR-induced mutagenesis

CRISPR mutagenesis was performed using the Alt-R^TM^ CRISPR-Cas9 System from Integrated DNA Technologies (IDT). For each gene, gRNAs were generated by mixing the crRNA and tracrRNA in a 1:1 ratio, diluting to 20 µM in nuclease-free duplex buffer (IDT), incubating at 95°C for 5 minutes and cooling on ice. Cas9 protein (20 µM, NEB) was mixed in a 1:1 ratio with the gRNA complex and incubated for 5-10 minutes at room temperate to produce the Cas9:gRNA RNP complex at a final concentration of 10 µM. RNPs were loaded into pre-pulled microcapillary needles (Tritech Research), calibrated, and 2 nL RNP complexes were injected into the yolk of 1-to 4-cell stage embryos.

### Sequencing and mutation efficiency analysis

Between 24 and 96 hpf, a few embryos from each injection group were pooled, DNA extracted, Sanger sequenced (GenScript), and screened for editing efficiency using the TIDE webtool (50). Individual animals were also screened for mutagenesis using whole larvae at 12 dpf to predict editing efficiencies using the TIDE webtool, and to check for clonal fitness effects. These data are reported as intra-animal efficiencies in the main text. For next generation sequencing, a single *wnt16* sample was prepared according to the recommended guidelines for CRISPR amplicon sequencing at the MGH CCIB DNA Core affiliated with Harvard University. High-quality reads were aligned, identical and similar sequences (those with isolated SNPs not affecting indels were considered similar) were collapsed, and counts of unique alleles with more than 10 paired reads are reported.

### MicroCT scanning and image analysis

MicroCT scanning was performed using a vivaCT40 (Scanco Medical, Switzerland) with 21 μm isotropic voxel resolution, and analyzed using FishCuT software (1, 32).

### Fluorescent imaging

Between 10-12dpf, zebrafish of the transgenic *sp7:EGFP* (23) background were anesthetized in MS-222 and mounted into borosilicate glass capillaries using 0.75% low melt-agarose (Bio-Rad) diluted in system water containing 0.01% MS-222. Dual-channel (GFP, excitation 450-490, emission 500-550; DAPI, excitation 335-383, emission 420-470) images were collected on a high-content fluorescent microscopy system (Zeiss Axio Imager M2, constant exposure settings for all experiments) using a 2.5× objective (EC Plan-Neofluar 2.5×/0.075). Maximum intensity projections were generated from image stacks in Fiji (51) for analysis. Of note, grouped breeding of this line yields ~95% positive “glower” fish (when screened at 24hpf) suggesting breeders and G0 crispants from this line are comprised of at least 50% of fish homozygous for the transgene. Notably, Eq 1 is expected to hold for clones representing mono-or multi-allelic loss, and thus our use of animals with mixed transgenic zygosities is not expected to influence our results. G0 crispants used for assessment of clonal dynamics were excluded if no distinct fluorescence could be detected above background during imaging. No attempt was made to quantify relative levels of fluorescence within or between fish. Further, loss of fluorescence (due to gene editing of either one copy or multiple copies of the transgene) was defined as complete loss of fluorescence (i.e. signal was indistinguishable from background).

### Monte Carlo simulations

We previously characterized multivariate distributions for select FishCuT measures that exhibit evidence of multivariate normality (1). Using parameter estimates previously derived for such measures, we constructed wildtype and mutant distributions using methods described in Supplemental Information. Mutant phenotypic distributions were parameterized by several variables: d (characteristic effect size), λ (extent of intra-animal variation in phenotypic expressivity; λ=0 is most variable, λ=1 is least variable), and p_i_ (a ‘loss-of-function vector’ whose values range from 0 to 1, and which encodes the spatial pattern of loss-of-function in each vertebra). We examined two classes of loss-of-function vectors. The first class, 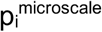, simulated phenotypes arising from microscale loss-of-function clusters. The second class, 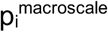, simulated macroscale loss-of-function clusters. When λ=1, mutant distributions were identical for both microscale and macroscale clusters; because loss-of-function was uniform in this case, we refer to simulated mutants with λ=1 as “germline mutants”. For simulations, unless otherwise noted, we assumed a characteristic effect size of d=2, and a sample size of n=10, or a characteristic effect size of d=2.5, and a sample size of n=15. Similar methods were used for Monte Carlo simulations for Moran’s I.

### Statistical analysis

In general, the Mann-Whitney test was used for univariate analyses between two groups. Dysmorphic phenotypes were analyzed using a test for equal proportions. Multivariate analysis was performed using the global test. p<0.05 was considered statistically significant in all cases.

## ACKNOWLEDGEMENTS

Research reported in this publication was supported by the National Institute of Arthritis and Musculoskeletal and Skin Diseases (NIAMS) of the National Institutes of Health (NIH) under Award Number AR066061 and AR072199. The content is solely the responsibility of the authors and does not necessarily represent the official views of the National Institutes of Health. ZIRC is supported by NIH grant RR12546. We thank Drs. Peter Byers and Cecilia Moens for helpful discussions.

---

**Fig 5-Supplemental Fig 1.**
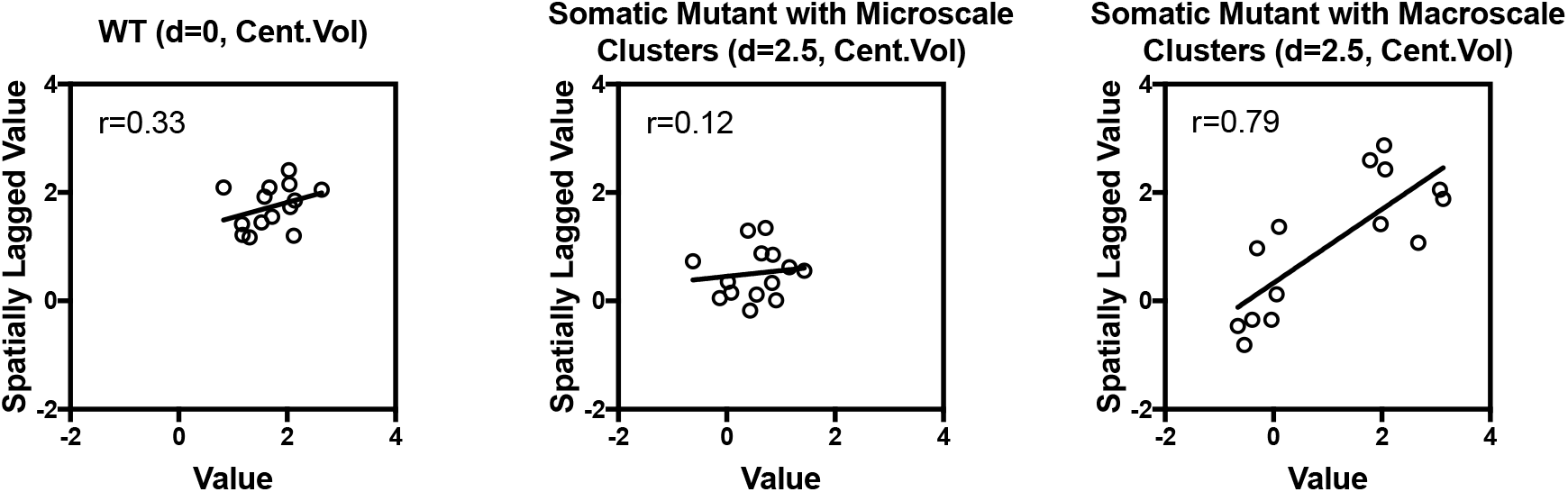
Numerical basis for changes in Moran’s I in the presence of microscale or macroscale loss-of-function clusters. Shown are Moran’s scatter plots, which contain the original variable (e.g. Cent.Vol in vertebra 4) on the x-axis in individual fish, and the spatially lagged variable (e.g. the mean of Cent.Vol in vertebrae 3 and 5) on the y-axis. For wildtype animals (left), there is natural phenotypic co-variation between vertebrae, resulting in a moderate spatial correlation. For somatic mutants with microscale clusters (middle), there is a greater prevalence of vertebrae with high phenotypic severity adjacent to those with low severity (and vice versa), resulting in loss of correlation in neighboring vertebrae. For somatic mutants with macroscale clusters (right), vertebrae within and outside of the cluster separate into the upper right and lower left quadrant, respectively, resulting in increased spatial autocorrelation in each animal.

**Fig 6-Supplemental Fig 1.**
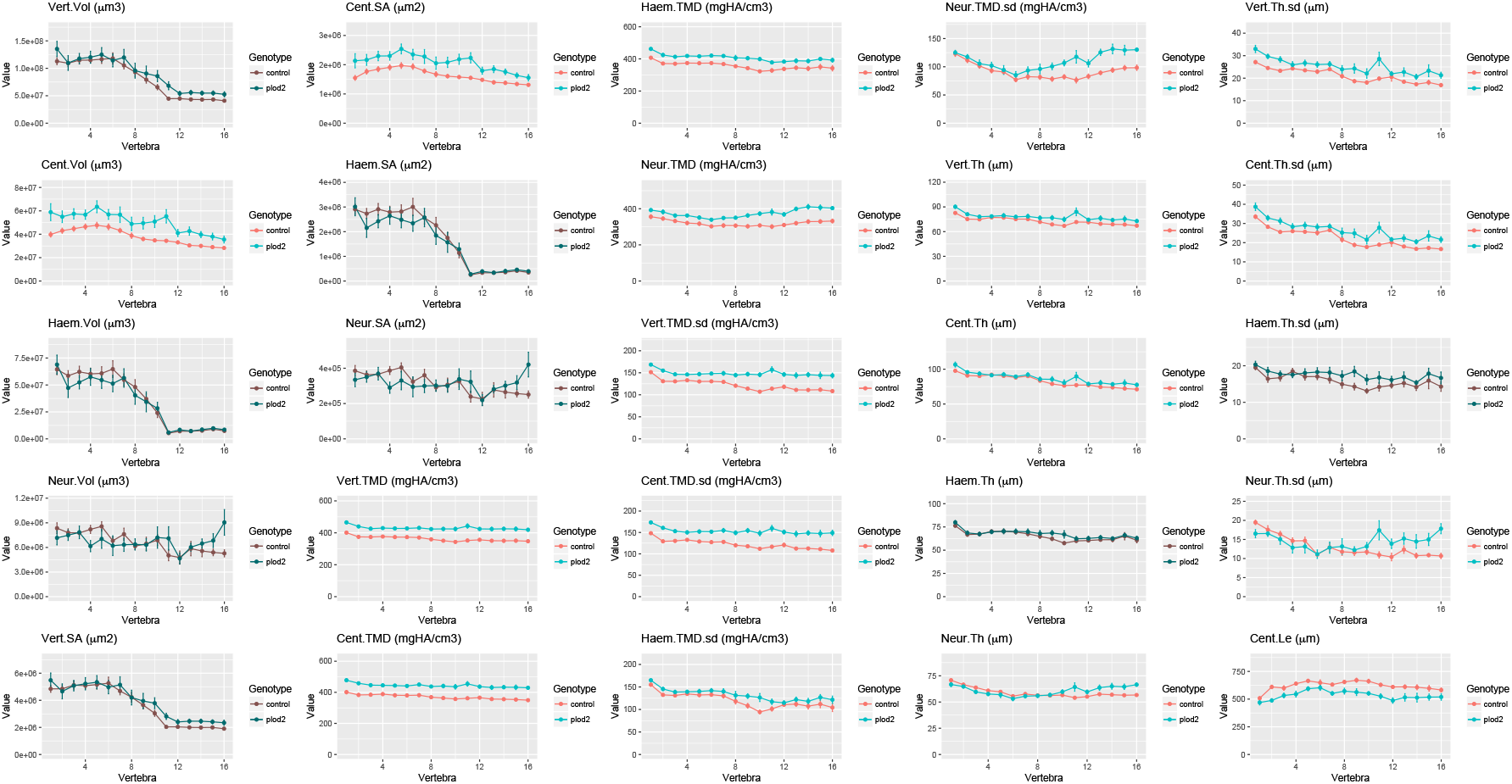
FishCuT analysis of *plod2* somatic mutants for all 25 combinatorial measures. Phenotypic features plotted as a function of vertebra (mean ± SE, n = 11/group). Plots associated with p<0.05 in the global test are colored in a lighter coloring scheme.

**Fig 7-Supplemental Fig 1.**
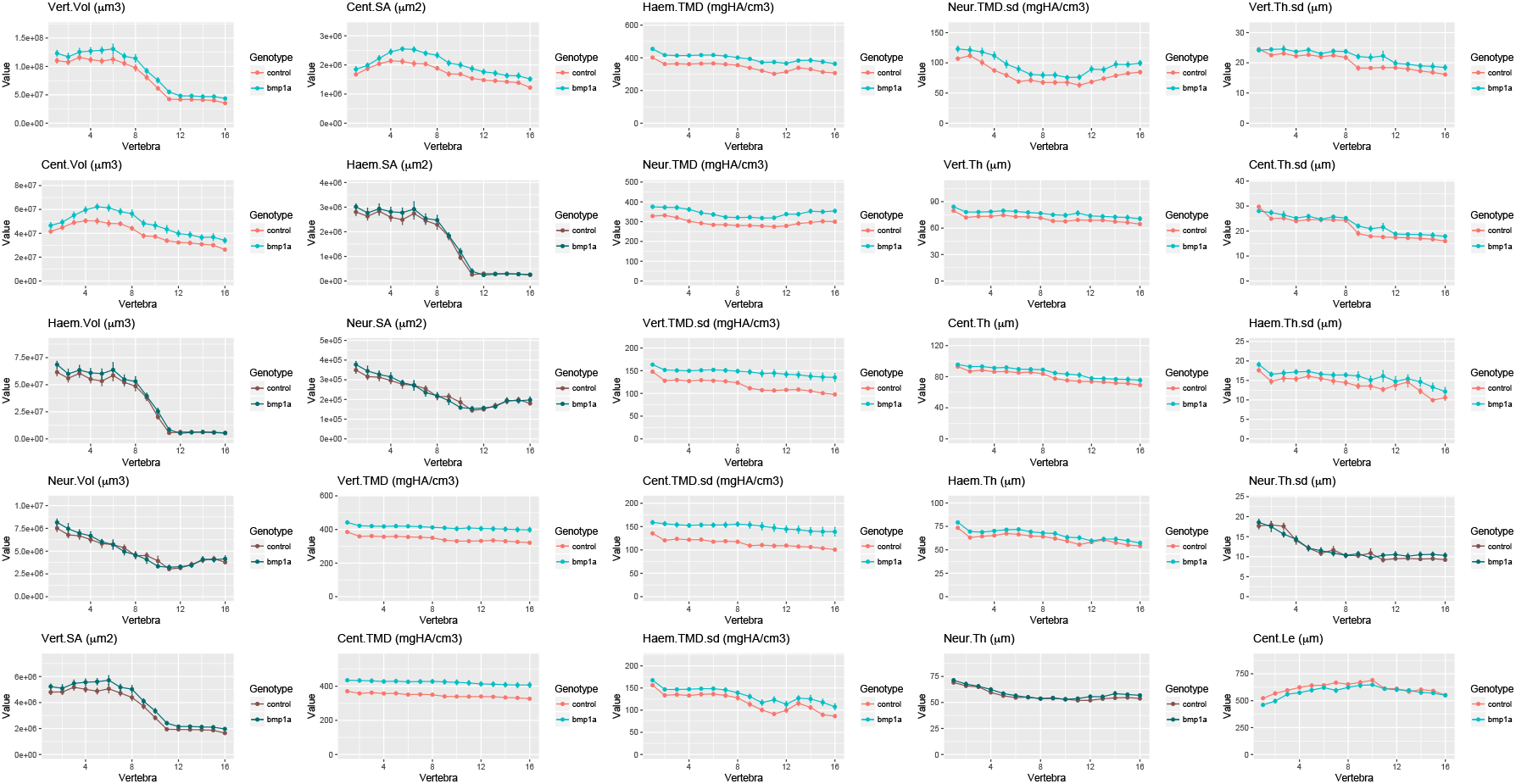
FishCuT analysis of *bmp1a* somatic mutants for all 25 combinatorial measures. Phenotypic features plotted as a function of vertebra (mean ± SE, n = 15/group). Plots associated with p<0.05 in the global test are colored in a lighter coloring scheme.

**Fig 8-Supplemental Fig 1.**
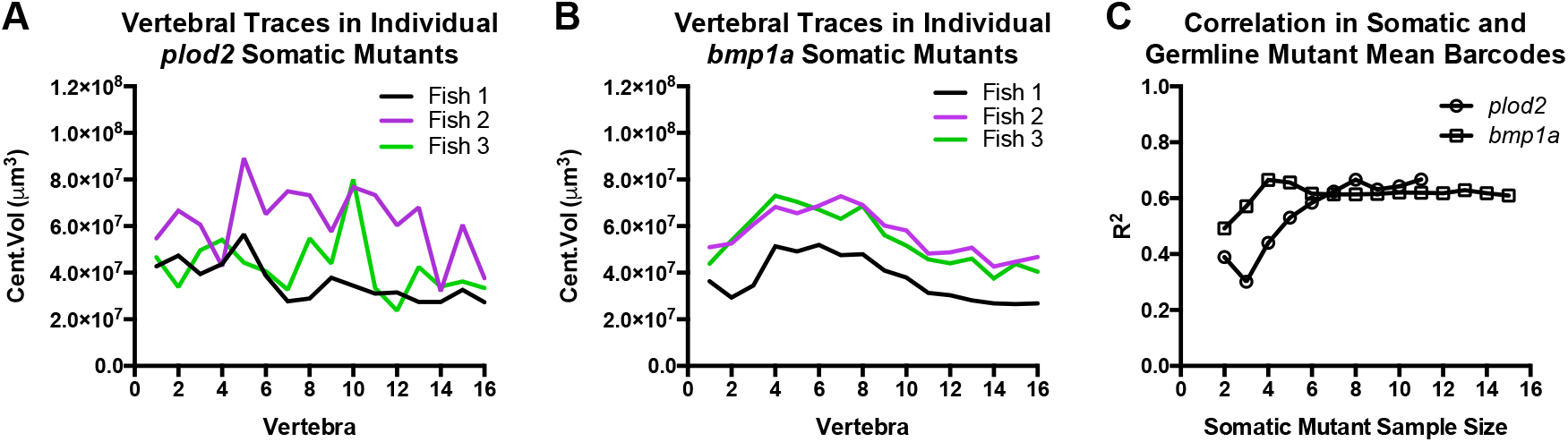
Somatic mutants for *bmp1a* and *plod2* exhibit inter- and intra-animal phenotypic variability that becomes averaged out across the sample. (A-B) Vertebral traces for Cent.Vol for individual *plod2* (A) and *bmp1a* (B) somatic mutants. Traces for *plod2* somatic mutants are jagged, whereas those for *bmp1a* somatic mutants are smooth. Thus, *plod2* and *bmp1a* somatic mutant groups exhibit different degrees of spatial phenotypic variability. Both somatic mutants groups exhibit variability in expressivity across individuals. (C) Correlation in somatic and germline mutant mean barcodes as a function of somatic mutant sample size. As sample size increases, phenome-wide correlation increases in an asymptotic manner. Minimal increases in correlation are observed beyond a sample size of n=8.

**Fig 9-Supplemental Fig 1.**
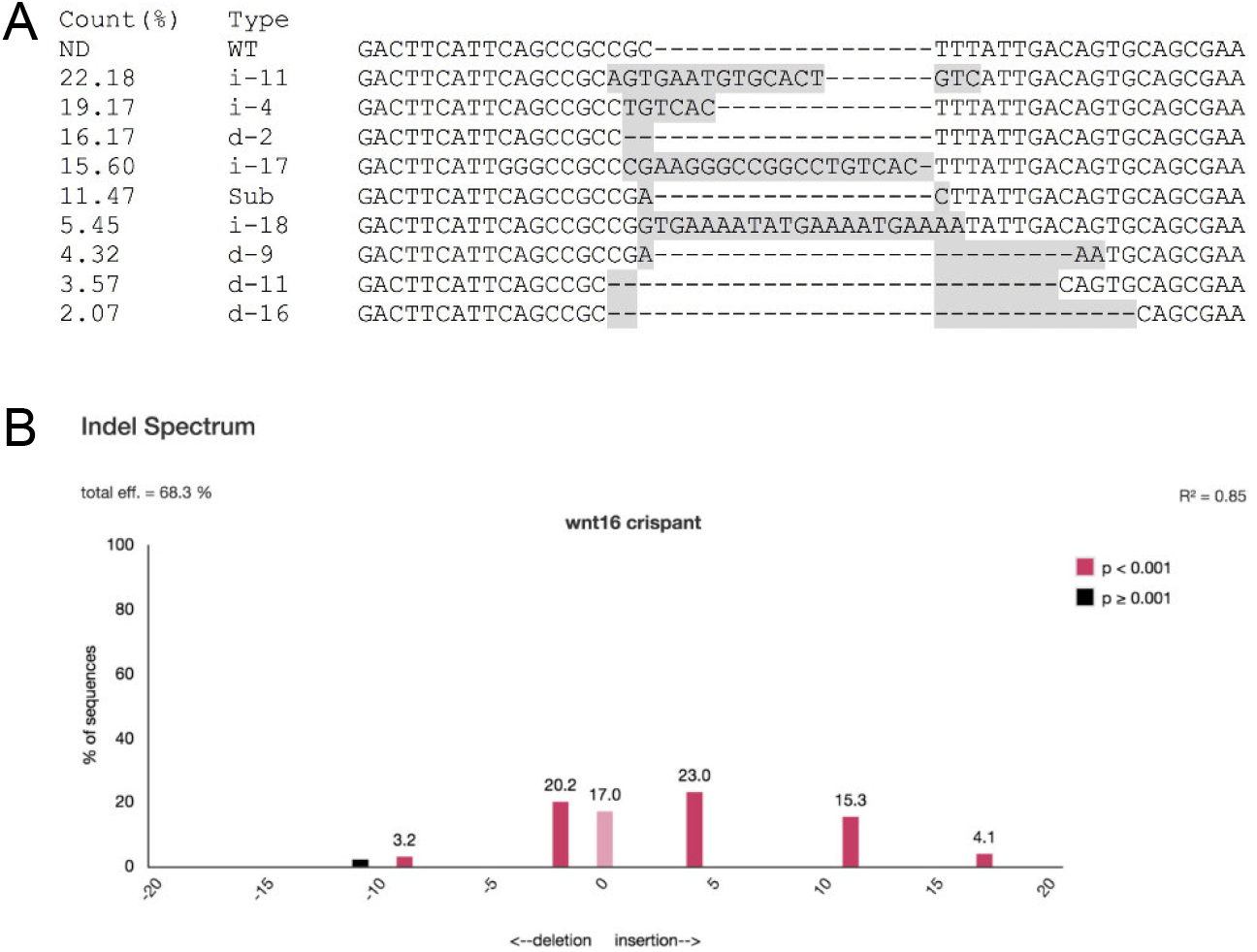
Indels in *wnt16* somatic mutants. (A) NGS data of a single 12dpf *wnt16* somatic crispant, showing the predominance of mutated alleles (represented as a % of total high quality NGS reads), allele type and sequence. Gray sequences show difference from wild type. i, insert; d, deletion; ND, not detected; Sub, substitution. (B) TIDE prediction for NGS sample in “A” showing relatively high correspondence in predicted alleles.

**Fig 10-Supplemental Fig 1.**
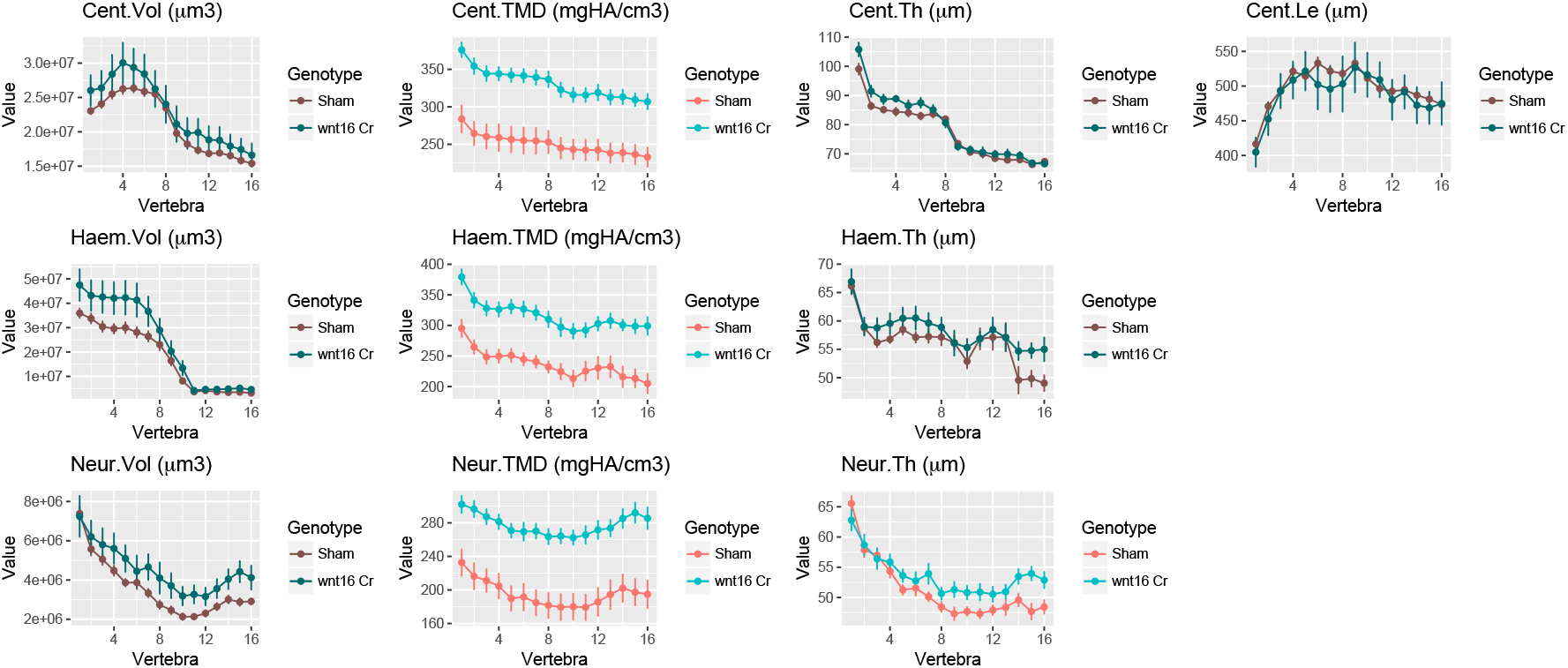
FishCuT analyses of *wnt16* somatic mutants with allometric normalization. Phenotypic features are plotted as a function of vertebra (mean ± SE, n=12/group). Plots associated with p<0.05 in the global test are colored in a lighter coloring scheme.

**Fig 10-Supplemental Fig 2.**
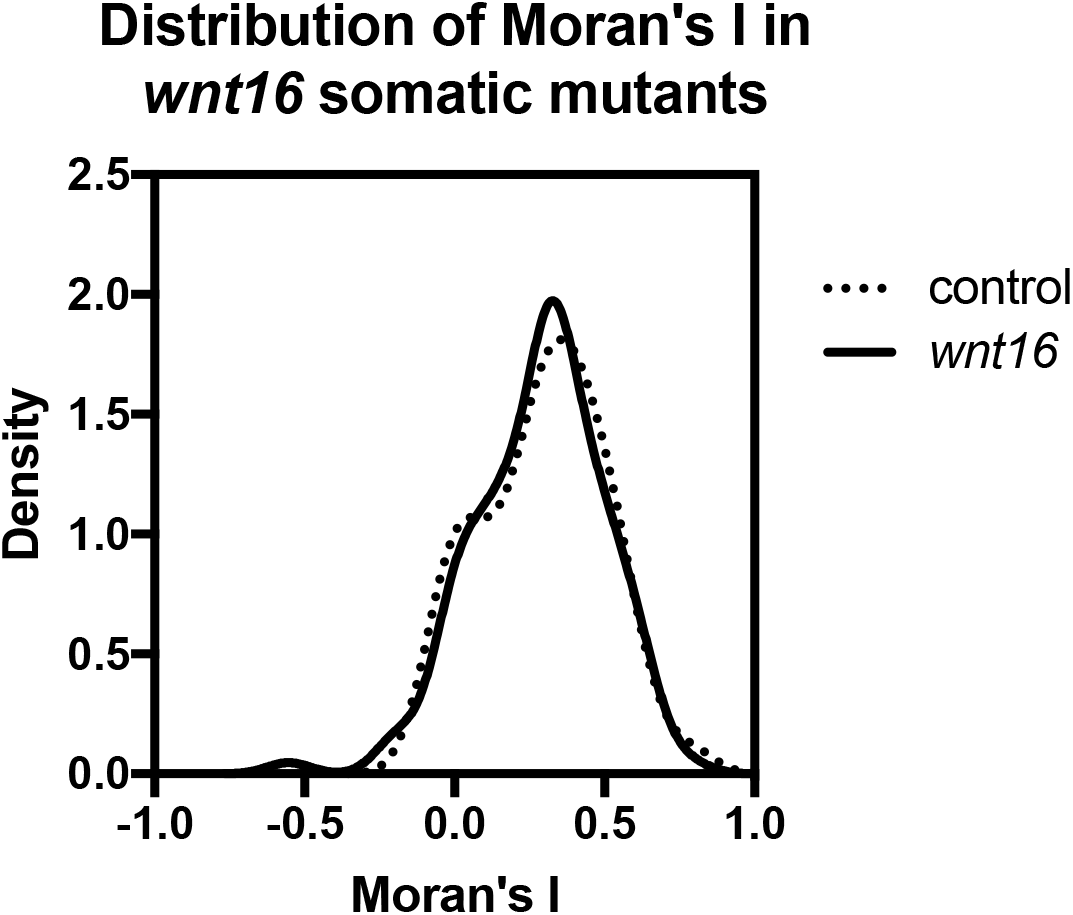
Distribution of Moran’s I in *wnt16* somatic mutants compared to controls.

**Fig 10-Supplemental Fig 3.**
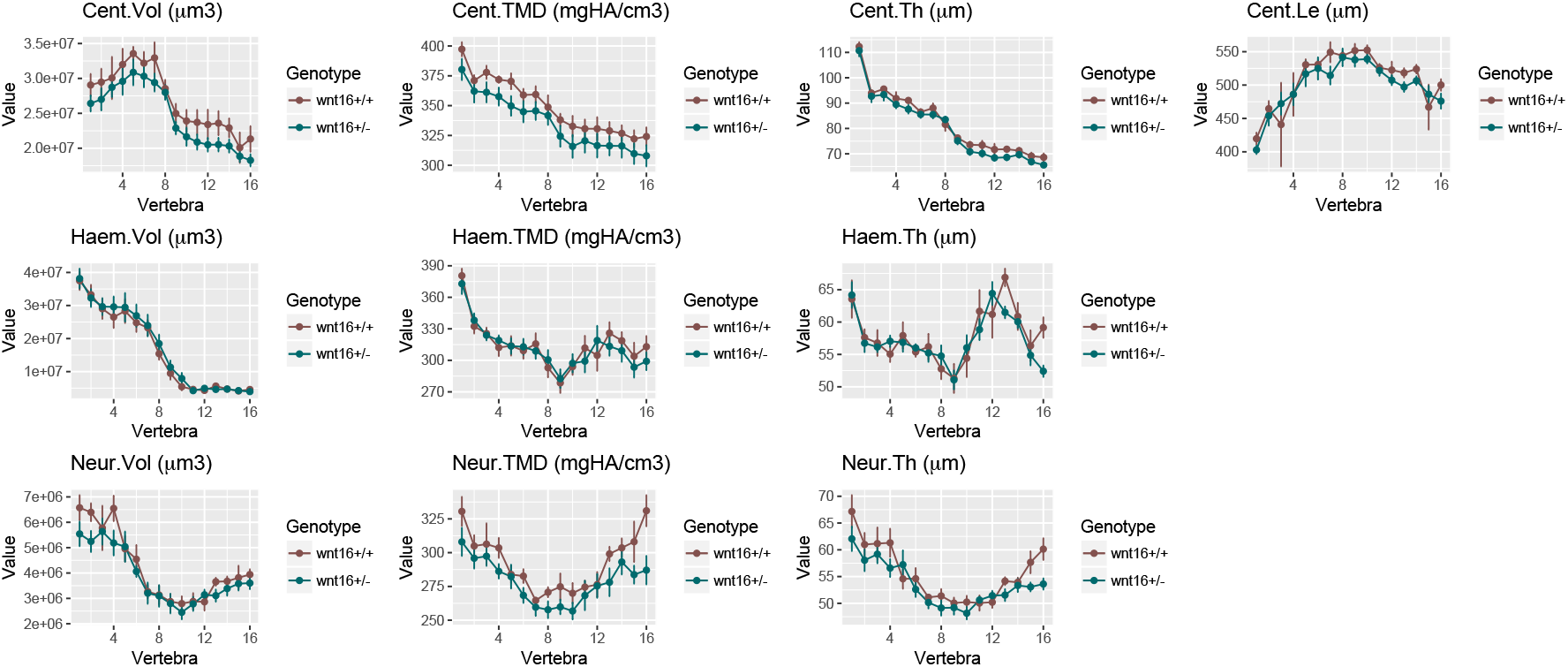
FishCuT analyses of *wnt16* heterozygous mutants. Phenotypic features are plotted as a function of vertebra (mean ± SE, +/−: n=6/group, +/+: n=4/group). Plots associated with p<0.05 in the global test are colored in a lighter coloring scheme.

