## Supplemental Information for "Decoding G0 somatic mutants through deep phenotyping and mosaic pattern analysis in the zebrafish skeleton"

Running Title: Decoding Somatic Phenotypes

Claire J. Watson[1,2,7], Adrian T. Monstad-Rios[1,2], Rehaan M. Bhimani[1,2], Charlotte Gistelinck[1,4], Andy Willaert[4], Paul Coucke[4], Yi-Hsiang Hsu[5,6], Ronald Y. Kwon[1,2,3]

[1] Department of Orthopaedics and Sports Medicine, University of Washington, Seattle, Washington, USA

[2] Institute for Stem Cell and Regenerative Medicine, University of Washington, Seattle, Washington, USA

[3] Department of Mechanical Engineering, University of Washington, Seattle, Washington, USA

[4] Center for Medical Genetics Ghent, Ghent University, Ghent, Belgium

[5] Hebrew SeniorLife Institute for Aging Research, Boston, Massachusetts, USA

[6] Harvard Medical School, Boston, Massachusetts, USA

Key Words: CRISPR, crispant, zebrafish, bone, osteoblast, phenomics, Osteogenesis Imperfecta, brittle bone disease, osteoporosis, *bmp1a*, *plod2*, *wnt16*

SUPPLEMENTARY TEXT

### DETAILED MATERIALS AND METHODS

#### *Zebrafish rearing*

All studies were performed on an approved protocol in accordance with the University of Washington Institutional Animal Care and Use Committee (IACUC). Zebrafish rearing was as described previously (1). Studies were conducted in mixed sex animals from either the wild-type AB strain or the transgenic *sp7:EGFP* (2) background. Stocks of both strains were obtained from the Zebrafish International Resource Center (ZIRC, http://zebrafish.org), and have been maintained for several generations in our facility. At the desired time point, zebrafish were euthanized by immersion in ice water, and stored frozen at -20°C. For all mutant lines, heterozygous mutant zebrafish were intercrossed to obtain homozygous mutants. All fish were housed in plastic tanks on a commercial recirculating aquaculture system.

#### *CRISPR-induced mutagenesis*

CRISPR mutagenesis was performed using the Alt-R™ CRISPR-Cas9 System from Integrated DNA Technologies (IDT). Target sequences were identified using the web-based tool, CHOPCHOP (3, 4), were designed using the GRCz10 reference genome (http://www.ensembl.org) and are as follows (PAM sequence in lowercase letters): `GATGGCCGCGTCGATTCTGGagg` for *sp7*, `GACCAGGATGGGCACCACCCcgg` for *EGFP*, `AAGTATCCGTCTGTACGCAGtgg` for *plod2*, `ATACGTGGGCCGCAGAGGAGggg` for *bmp1a*, and `CTGCACTGTCAATAAAGCGGcgg` for *wnt16.* The stable *wnt16* line harbored a 10 base pair insertion in exon 3 of the gene (reference sequence/mutant allele: `TTCAGCCGCCGCT[TT-AT/GTCATTTATTTAAA]TGACAGTGCAGC`). CRISPR-Cas9 crRNAs (gene-specific) and tracrRNAs (generic) were chemically synthesized and ordered from IDT. For each gene, gRNAs were generated by mixing the crRNA and tracrRNA in a 1:1 ratio, diluting to 20 µM in nuclease-free duplex buffer (IDT), incubating at 95°C for 5 minutes and cooling on ice. Cas9 protein (20 µM, NEB) was mixed in a 1:1 ratio with the gRNA complex and incubated for 5-10 minutes at room temperate to produce the Cas9:gRNA RNP complex at a final concentration of 10 µM. Sham complexes contained Cas9:tracrRNA at final concentration of 10 µM. RNPs were loaded into pre-pulled microcapillary needles (Tritech Research), calibrated, and 2 nL RNP complexes were injected into the yolk of 1- to 4-cell stage embryos.

*Sequencing and mutation efficiency analysis*

Between 24 and 96 hpf, a few embryos from each injection group were pooled, DNA extracted, Sanger sequenced (GenScript), and screened for mutagenesis. efficiency using the TIDE webtool (5). Individual animals were also screened for mutagenesis using whole larvae at 12 dpf to predict mutation efficiencies using the TIDE webtool, and to check for clonal fitness effects. These data are reported as intra-animal efficiencies in the main text. Furthermore, without exception for *bmp1a* and *plod2*, and in 9 of 12 (75%) individuals for *wnt16*, indel efficiencies were limited by the $R^2$ value in the TIDE analysis, suggesting that these may represent the lower boundary of CRISPR-induced indels for all samples and/or target sites. Only in 2 of 12 *wnt16* somatic mutants was a 0 bpd/bpi allele predicted to be represented by the sequence deconvolution, and one of these two samples was chosen for next-generation sequencing.

For next generation sequencing, a single *wnt16* sample was prepared according to the recommended guidelines for CRISPR amplicon sequencing at the MGH CCIB DNA Core affiliated with Harvard University. Raw fastq files retrieved from the MGH Core containing individual reads were analyzed using a combination of the Galaxy webtool (6), CLC Sequence Viewer (Qiagen) and Geneious software (Biomatters Ltd.). High-quality reads were to aligned, identical and similar sequences (those with isolated SNPs not affecting indels were considered similar) were collapsed, and counts of unique alleles with more than 10 paired reads are reported.

*MicroCT scanning and image analysis*

MicroCT scanning was performed using a vivaCT40 (Scanco Medical, Switzerland). Scans with 21 μm isotropic voxel resolution were acquired using the following settings: 55kVp, 145mA, 1024 samples, 500proj/180°, 200 ms integration time. DICOM files of individual fish were generated using Scanco software, and analyzed using FishCuT software (1, 7). Two fish were scanned simultaneously in each acquisition.

*Fluorescent imaging*

Between 10-12dpf, zebrafish were anesthetized in MS-222 and mounted into borosilicate glass capillaries using 0.75% low melt-agarose (Bio-Rad) diluted in system water containing 0.01% MS-222. Capillaries were set on a custom 3-D printed holder to aid manipulation and rapid orientation of the specimen. Dual-channel (GFP, excitation 450-490, emission 500-550; DAPI, excitation 335-383, emission 420-470) images were collected on a high-content fluorescent microscopy system (Zeiss Axio Imager M2, constant exposure settings for all experiments) using a 2.5x objective (EC Plan-Neofluar 2.5x/0.075). For each fish, a composite image stack (usually 3/1 images in the x/y directions; and optimized to 30-70 μm slice intervals in the z direction across the entire region of interest, usually about 9 slices; all at 2.58 μm/pixel) was acquired in mediolateral and anteroposterior views. Maximum intensity projections were generated from image stacks in Fiji (8) for analysis.

*Monte Carlo simulations*

We previously characterized multivariate distributions for select FishCuT measures that exhibit evidence of multivariate normality (1). Using parameter estimates previously derived for such measures, we constructed two distributions. The wildtype (WT) distribution, $y_i^{WT}$, consisted of a multivariate normal distribution using means $\mu_i^{WT}$ (which denotes the mean value in vertebra i in WT fish) and covariances $\Pi_{ij}^{WT}$ (which denotes the covariance between vertebra i and vertebra j in WT fish) (1). The mutant distribution was computed as

$$y_i^{mutant} = y_i^{WT} + d*\sigma_i^{WT} [(1-\lambda)p_i + \lambda] \quad \text{Eq 2}$$

where d is the characteristic effect size (1), $\sigma_i^{WT}$ is the standard deviation in vertebra i in WT fish, $\lambda$ is a parameter controlling intra-animal variability in loss-of-function that ranges from zero to one, and $p_i$ is a vector of values ranging from zero to one. When $\lambda$=0, Eq 2 simplifies to $y_i^{mutant} = y_i^{WT} + d*\sigma_i^{WT}*p_i$. Here, it can be seen that phenotypic severity varies from vertebra-to-vertebra, depending on values of $p_i$. We termed $p_i$ the 'loss-of-function vector' because it encodes the degree of phenotypic severity in each vertebra.

We constructed two classes of loss-of-function vectors. The first class, $p_i^{microscale}$, simulated phenotypes arising from microscale loss-of-function clusters. In this case, values of $p_i^{microscale}$ were drawn from the distribution of Eq 1, assuming $\langle x \rangle$=0.5 vertebrae. The second class, $p_i^{macroscale}$, simulated macroscale loss-of-function clusters. Here, each mutant possessed a single macroscale loss-of-function cluster whose size was drawn from the distribution of Eq 1. For this, we assumed $\langle x \rangle$=8 vertebrae, representing a moderately variable phenotype where half of the vertebrae, on average, are affected. For this class, $p_i$ was specified as a binary vector, with the center of the macroscale cluster drawn from a uniform distribution. In both cases, gene action was assumed to be cell autonomous, in that phenotypic severity in each vertebra was proportional to the extent of loss-of-function within it. Note that when $\lambda$=1, Eq 2 simplifies to $y_i^{mutant} = y_i^{WT} + d*\sigma_i^{WT}$, and is identical for both microscale and macroscale clusters. For simulations, unless otherwise noted, we assumed a characteristic effect size of d=2, and a sample size of n=10, or a characteristic effect size of d=2.5, and a sample size of n=15. All Monte Carlo simulations were performed in R (9).

*Moran's I*

Similar methods to those above were used for Monte Carlo simulations for Moran's I. Moran's I was computed using the Moran.I function in the ape package in R (9). The following weight matrix Wij was used:

Wij = 1 if abs(i-j)=1, else
Wij = 0

where i and j denote the $i^{th}$ and $j^{th}$ vertebra. We computed Moran's I for each vertebral trace as $I(d_i)$, where $d_i = [\underline{y}_i - \mu_i^{WT}]/\sigma_i^{WT}$, and $\underline{y}_i$ is a vertebral trace drawn from the distribution of Eq 2.

*Statistical analysis*

For mutant fish with a known axial skeletal phenotype (*bmp1a* and *plod2* somatic mutants), results are reported from a single experiment; for characterization of mosaicism in *sp7:EGFP* fish, results are reported from two experiments. For somatic *wnt16* mutants, results are reported from three experiments. Because germline *wnt16* mutants recapitulated somatic mutant phenotypes, results are reported from a single experiment. Each biological replicate represents one technical replicate. Mild and extreme outliers were identified only for descriptive purposes; all data were included in statistical analyses. All statistical analyses were performed in GraphPad Prism or R [25]. In general, the Mann-Whitney test was used for univariate analyses between two groups. Dysmorphic phenotypes were analyzed using a test for equal proportions. Multivariate analysis was performed using the globaltest package (10). $p<0.05$ was considered statistically significant in all cases.
